# How tactile afferents in the human fingerpad encode tangential torques associated with manipulation: are we better than monkeys?

**DOI:** 10.1101/2022.07.11.499647

**Authors:** Alastair J. Loutit, Heather E. Wheat, Heba Khamis, Richard M. Vickery, Vaughan G. Macefield, Ingvars Birznieks

## Abstract

Dexterous object manipulation depends critically on information about forces normal and tangential to the fingerpads, and also on torque associated with object orientation at grip surfaces. In this study we investigated how torque information is encoded by human tactile afferents, including slowly adapting type-II (SA-II) afferents, in the fingerpads. SA-II afferent properties seemed perfectly suited for torque encoding but could not be previously investigated as they are absent in the glabrous skin of monkeys. Torques of different magnitudes (3.5-7.5 mNm) were applied in clockwise and anticlockwise directions to a standard central site on the fingerpads of 34 participants. Torques were superimposed on a 2 N, 3 N, or 4 N background normal force. Unitary microneurography recordings were made from fast adapting type-I (FA-I, n=39), slowly adapting type-I (SA-I, n=31), and type-II (SA-II, n=13) afferents supplying the fingerpads. All three afferent types encoded torque magnitude and direction, with SA-II afferents showing excitatory and inhibitory modulation depending on torque direction. Most afferents of all three types had higher torque sensitivity with smaller normal force. FA-I afferents showed the best torque magnitude and worst directional discrimination abilities in both species. Human SA-I afferent response to static torque was inferior to dynamic stimuli, while in monkeys the opposite was true. In humans this might be compensated by the addition of a sustained SA-II afferent input. In comparison to monkeys the performance of each afferent type was inferior in humans, likely due to differences in fingertip tissue compliance and skin friction. [currently 246 words; 250 words max]

**Significance Statement:** We investigate how individual human tactile nerve fibres encode rotational forces (torques) and compare them to their monkey counterparts. Human hands, but not monkey hands, are innervated by a tactile neuron type (SA-II afferents) specialised to encode directional skin strain, yet so far, torque encoding has only been studied in monkeys. We find that human SA- I afferents were generally less sensitive and less able to discriminate torque magnitude and direction than their monkey counterparts, especially during the static phase of torque loading. However, this shortfall in humans could be compensated by the addition of SA-II afferent input. This indicates that variation in afferent types might compliment their specialisation for stimulus features, by slight differences between encoding in the two species.

## Introduction

When manipulating held objects, rotational forces (torques) commonly develop at the thumb and fingers. Sensing those forces might be important to control an object’s orientation in the hand. For example, increasing the grip force on a knife to counteract developing torque sensed by the fingertips when cutting. Additionally, greater grip force is required to prevent rotational than tangential slips, which can be exploited to reorient grasped objects using controlled torsional slips, like allowing a glass of water to rotate between our fingers to align with gravity and thereby prevent spills, without inducing net vertical translation of the contact area (tangential slip) (Johansson and Westling, 1984; Kinoshita et al., 1997). It follows that to achieve an action goal, the grip force should be precisely adjusted to both tangential forces and torque, whether it be to achieve a desired object orientation by maintaining constant torque levels and avoiding slips, or to purposefully exploit controlled rotational slips.

Skin mechanoreceptors, and their corresponding afferents, contribute to the control of dexterous object manipulation by signalling gripped object features, fingertip forces, and grip safety (Johansson and Westling, 1987; Goodwin et al., 1998; Nowak et al., 2003; Johansson and Flanagan, 2009; Crevecoeur et al., 2011; Khamis et al., 2014; Delhaye et al., 2021; Schiltz et al., 2021). Sensory inputs can rapidly update internal representations of hand-object manipulations, which are used to counteract unexpected events and build predictive strategies (Johansson and Westling, 1988; Johansson and Cole, 1992; Miall and Wolpert, 1996; Birznieks et al., 1998; Witney et al., 2004; Flanagan et al., 2008) which might also include planned slips (Birznieks et al., 1998).

Afferent torque responses have previously been characterised only in macaques. Most fast-adapting type I (FA-I) and slowly-adapting type I (SA-I) low-threshold mechanoreceptive afferents, innervating the fingertip, had response impulse rates that were scaled by torque magnitude (Birznieks et al., 2010; Redmond et al., 2010a; Redmond et al., 2010b; Khamis et al., 2015). SA-I afferents showed preference for torque direction, whereas FA-I afferents responded to torque magnitude, irrespective of direction. Models trained to decode afferent input demonstrated that several combinations of normal force, torque magnitude, and direction could be accurately classified (Birznieks et al., 2010; Redmond et al., 2010a; Redmond et al., 2010b; Fu et al., 2012), and concurrently extract instantaneous grip force and torque parameters from a small number of tactile afferents’ responses, in a real-time fashion (Khamis et al., 2015).

Until now, no data from human tactile afferents have been available for such modelling. Unlike monkeys, human glabrous skin possesses a prominent population of slowly-adapting type II (SA-II) afferents showing exquisite sensitivity to the direction and magnitude of skin stretch (Knibestöl and Vallbo, 1970; Knibestöl, 1975; Johansson, 1978; Olausson et al., 2000; Birznieks et al., 2001; Birznieks et al., 2009), and eliciting sustained percepts of diffuse pressure, squeezing, or strain (Macefield et al., 1990; Kunesch et al., 1995; Watkins et al., 2022), thus making them ideal candidates for sensing torque. Moreover, biomechanical differences in fingertip structures may influence receptor response properties to stimuli involving gross skin deformation, such as torques which develop over the skin-object contact area. Human fingertips are larger and flatter than most nonhuman primate species, and differ in viscoelastic and frictional skin properties (Dandekar et al., 2003), which could lead to torque-encoding differences among species.

Here, we quantify human single FA-I, SA-I and SA-II afferent torque responses, recorded in the median nerve; the stimuli were too slow to activate FA-II (Pacinian) afferents. We were particularly interested in the potential signalling advantages offered by SA-II afferents, which are present in human, but not monkey, hand glabrous skin (Johnson, 2001). We applied clockwise and anticlockwise torques of three different magnitudes, to a central site on the glabrous skin of human distal fingertips, superimposed on three different background normal forces. First, we illustrate and characterise afferent response properties to torque, then we compare the torque discriminative capacity we observed in human and monkey afferents.

## Materials and methods

### Single-unit tactile afferent recordings

Single-unit tactile afferent recordings were acquired from 34 participants (median age 22, range 19-54; 19 females) using standard microneurography procedures (Vallbo and Hagbarth, 1968; Ackerley and Watkins, 2022). Each participant provided informed written consent to the procedures in accordance with the Declaration of Helsinki, and all procedures were approved by the UNSW Human Research Ethics Committee. Participants reclined in a dental chair with their forearm supinated and Velcro straps around the wrist were used to secure the forearm to a vacuum cast to immobilise the arm. The dorsum of the hand was embedded in plasticine up to the mid-level with the fingers splayed but the distal end of the phalanges did not contact the plasticine, thereby allowing normal skin deformation mechanics during mechanical stimulation. To stabilize the digits, the nails of the index, middle, and ring fingers were glued to metal posts firmly sunk into the plasticine, and the thumb and little finger were secured by ‘U’ shaped aluminium clamps embedded in the plasticine.

Electrical impulses were recorded from individual low-threshold mechanoreceptive afferents with cutaneous receptive fields in the distal phalanx of the index, middle, or ring finger by percutaneously inserting tungsten needle electrodes (FHC Inc, Bowdoin, ME, USA) into the median nerve at the wrist. Neural signals were amplified (20,000 X) and filtered (300 Hz – 5 kHz) through an isolated amplifier (ISO-Dam 80, WPI, Sarasota, FL, USA). The microelectrode was manually guided into a cutaneous fascicle by delivering weak electrical pulses (0.2 ms, 1 Hz, 0.01-1.00 mA) via an isolated stimulator (Stimulus Isolator, ADInstruments, Sydney, NSW, Australia); radiating paraesthesia at 0.02 mA indicated that the tip had impaled a fascicle. Once the fascicle had been identified, the microelectrode was manipulated while the distal phalanx was mechanically stimulated and action potentials from a single sensory axon encountered.

### Classifying the afferent population

Calibrated nylon filaments (Semmes-Weinstein aesthesiometers, Stoelting, Chicago, IL, USA) were used to probe the receptive field borders of isolated afferents, defined as the region of skin from which a response could be elicited by a force four times the threshold force at the most sensitive zone in the receptive field. If an afferent was spontaneously active, the smallest force that could be seen to modulate its discharge rate (typically by about 10%) was defined as the threshold force.

Afferents were classified as fast adapting (FA) or slowly adapting (SA) by their responses to sustained indentation, and as type I or type II if they had small discrete receptive fields or large receptive fields with poorly defined borders, respectively. To differentiate SA-I and SA-II afferent responses, we also determined whether afferents fired spontaneously, the sharpness of the discharge rate increase at stimulus onset, and the variability of the discharge rates during the plateau phases of force and torque stimuli. To evaluate the discharge variability of SA-I and SA-II afferents during the force and torque plateau phases, we used an irregularity index. The irregularity index was calculated as the mean of the absolute discharge rate difference measured between every two pairs of consecutive spikes divided by the mean discharge rate (Birznieks et al., 2008).

We recorded from 83 low-threshold mechanoreceptive afferents with receptive field centres (RFCs) on the fingertip glabrous skin of digits 2, 3, and 4 of the left hand. Thirty-nine afferents were classified as fast adapting type I (FA-I), thirty-one afferents were classified as slowly adapting type one (SA-I), and thirteen as slowly adapting type two (SA-II). We excluded recordings from FA-II afferents because they did not respond reliably to our stimuli (Birznieks et al., 2001; Birznieks et al., 2010).

Previously published neural spike data acquired from 25 FA-I and 58 SA-I afferents in three anaesthetised *Macaca nemestria* monkeys, as described in Birznieks et al. (2010), were also used to compare human and monkey afferent torque responses.

### Stimulus

A custom-made stimulator was used to deliver normal forces and torques. This is the same stimulator as used for the studies in macaques (Birznieks et al., 2010). The stimulus applicator was a disc (24 mm diameter) with a flat stimulation surface. The disc was positioned so that the rotational torque axis was aligned over the centre of the flat portion of the volar surface of the fingertip and perpendicular to the skin surface (**Figure 1A**). This location remained the standard test site on the receptor-bearing finger, referred to as the rotational centre, regardless of receptor location.

**Figure 1.**
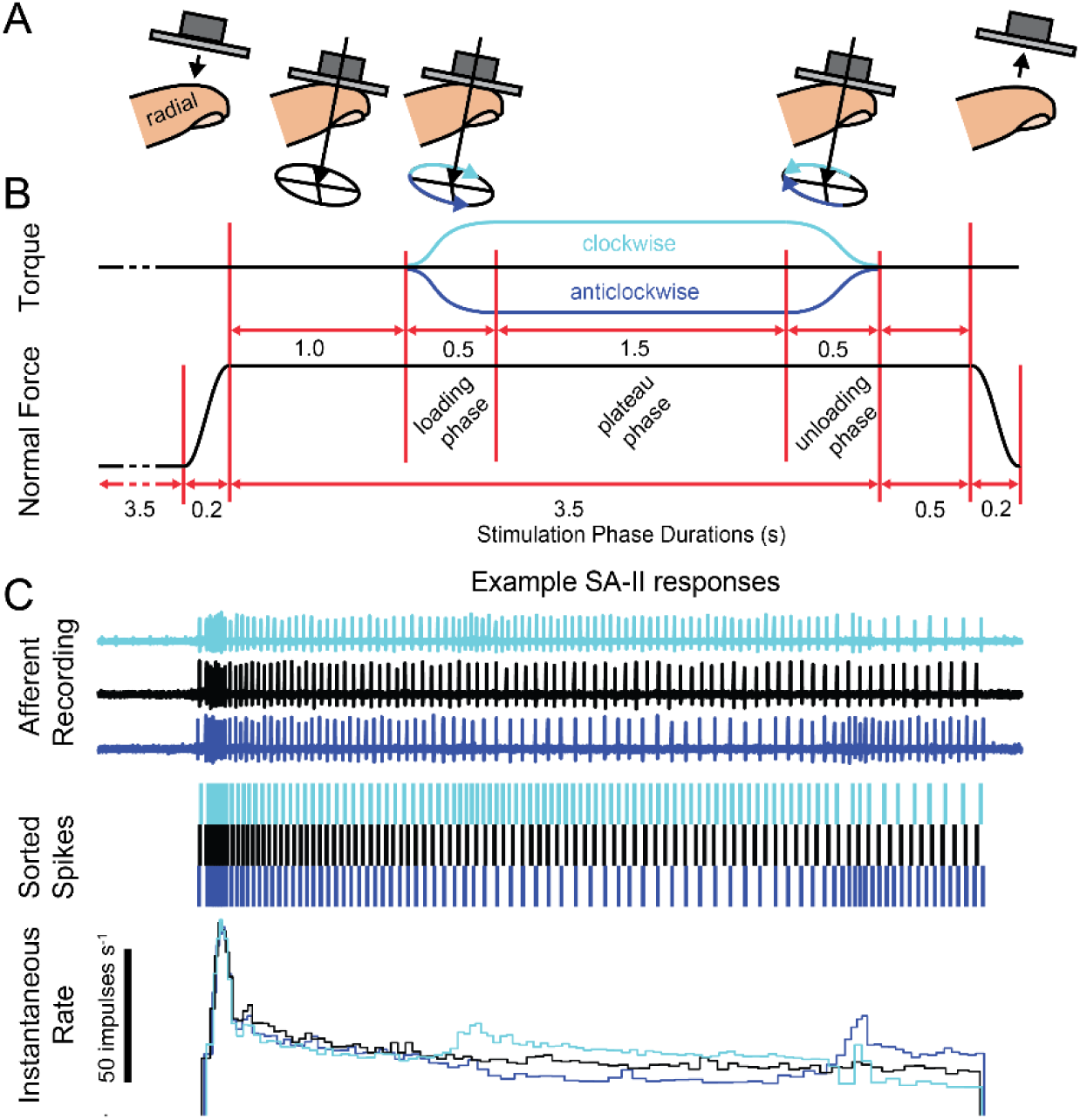
Normal force and torque stimulus application phases and an example of a single SA-II afferent response. **A**) Schematic representation of the stimulus applicator during fingertip stimulation. The surface was oriented parallel to the flat portion of the fingertip and was held just above the skin surface. The centre of the rotational axis for torque applications is indicated by black arrows. In each trial, the stimulator was advanced to compress the skin until the desired normal force was reached, then torque was applied, held at the plateau then it was rotated back to the starting point after which the applicator retracted from the skin surface to return to zero normal force. **B**) The phases of normal force and torque applications over time. Different magnitudes of torques were tested in clockwise (cyan) and anticlockwise (blue) directions. **C**) Examples of an SA-II afferent recording in response to normal force only (black) and normal force and torque applications in the clockwise (cyan) and anticlockwise (blue) directions. Inspected and sorted spikes are shown in the middle panel. The bottom panel shows the corresponding instantaneous rates.

Torques were applied at 3.5 mNm, 5.5 mNm, and 7.5 mNm in clockwise and anticlockwise directions. Each of the torque magnitudes and directions were applied with three different normal forces: 2 N, 3 N, and 4 N, corresponding to grip forces generated during object manipulation. Forces and torques applied to the fingertip were measured by a six-axis, force-transducer (Nano F/T; ATI Industrial Automation, Apex, NC) at 1000 Hz sampling rate with a force resolution of 0.0125 N and a torque resolution of 0.0625 mNm. The stimulus applicator was lowered to slightly above the skin and the damper setting adjusted to facilitate smooth contact with the skin surface. Each trial consisted of a dynamic phase of increasing normal force for 0.2 s, a static phase in which normal force was constant for 4.0 s, and a dynamic retraction phase lasting 0.2 s. Torques were superimposed during the constant normal force phase, commencing 1 s after the beginning of this phase (**Figure 1B**). The torque loading phase lasted 0.5 s, was held constant for 1.5 s during a plateau phase, followed by an unloading phase which lasted 0.5 s. Normal force unloading commenced 0.5 s after the end of the torque unloading phase (**Figure 1B**). Example traces from an SA-II afferent responding to each of the torque phases, and the neural spikes sorted for analysis are shown in **Figure 1C**. An example of the calculated instantaneous discharge rate is shown at the bottom.

Each stimulus set with a given background normal force consisted of presenting two normal- force only trials followed by three trials, each with one of the three torque magnitudes (3.5, 5.5, and 7.5 mNm) superimposed in ascending order. One experimental run comprised this stimulus set repeated three times, which took approximately two minutes. An experimental run was then repeated, resulting in six recordings of responses to stimuli of each torque magnitude and direction. Sometimes, when recording quality and time permitted, additional experimental runs were obtained, resulting in nine or twelve recordings for each torque magnitude and direction. The experimental runs were presented with 2 N normal force and in one torque direction and then the two runs were repeated for the same normal force but in the opposite torque direction. The full procedure was repeated with 3 N and 4 N normal forces. In most cases torques superimposed on 2 N normal force were tested before 3 N and 4 N, so there was a larger pool of data with 2 N normal force. Together, there were eighteen normal force and torque combinations: three torques presented in each direction (n=6), each superimposed on three normal forces. Presentations of each normal force torque combination were typically repeated six times. However, due to limited recording time in human studies, as signal-to-noise ratio sometimes declined before all stimuli could be tested not every afferent could be tested with all eighteen normal force and torque combinations and the desired number of repeats.

### Standardised finger

To standardise data collected from different subjects and fingers, receptive field centres RFCs were referenced to a standardised fingertip on the right hand, by normalising the coordinates of each fingertip and the RFC distances from the stimulus centre to the measurements of the average fingertip size. Afferent RFCs on the left hand and torque directions applied to the left hand were treated as mirror images of the right hand and plotted accordingly on the standardised finger (Birznieks et al., 2010). Therefore, clockwise was defined as rotation of the stimulus applicator from ulnar to distal to radial aspects of the fingertip, irrespective of the hand stimulated.

### Signal analysis

Afferent neural activity was acquired using a PowerLab 16/35 data acquisition system and viewed in LabChart 5 (ADInstruments, Sydney, NSW, Australia). Recorded neural spikes were sorted and selected for further analysis using custom-written code in Igor Pro 5 (Wavemetrics, Portland, OR, USA). Sorted spikes were analysed using custom-written code in MATLAB (The Mathworks, Natick, MA, USA). In response to each stimulus, we calculated the mean discharge rate in the torque loading phase, plateau phase, and unloading phase. We also calculated the instantaneous peak discharge rate during the dynamic loading and unloading phases. For afferents without ongoing activity (generally FA-I and SA-I) we also quantified the latency of occurrence of the first spike after the beginning of the torque loading phase.

### Data analysis

To determine whether afferent discharge rates were correlated with torque magnitudes, we calculated Spearman’s rank correlation coefficients between the mean discharge rate and torque magnitude. Correlation coefficients were calculated separately for each torque direction and each normal force magnitude. Responses to normal force only (zero torque) were not included in the correlation analysis. Afferents that responded to torque but did not show response scaling to the torque magnitudes tested in this study were analysed to determine if they had a non-graded response to torque by comparing responses of normal force only trials to trials with torque applications, regardless of magnitude, using the Mann-Whitney U test (MATLAB, The Mathworks, Natick, MA, USA). To compare torque effects between afferents and between stimulus conditions, we calculated torque sensitivity over three torque ranges: low torques 0-3.5 mNm, *S*_T_^low^; high torques 3.5-7.5 mNm, *S*_T_^high^; and the whole torque range 0-7.5 mNm, *S*_T_^all^. The number of spikes per second was plotted against the torque magnitude, and torque sensitivities were calculated as the gradient of the regression line. Thus,

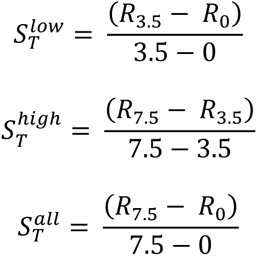

where R*_Z_* is the number of spikes at torque magnitude *Z*, and the resulting sensitivity was the number of spikes per mNm of torque per second. When torque sensitivity analysis was applied to the afferent population we used the absolute sensitivity values, 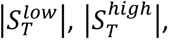 and 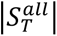. To compare how much afferent responses were influenced by the same amount of torque changes within low versus high range of torque, we calculated a torque sensitivity increase index, *I*_T,_ as

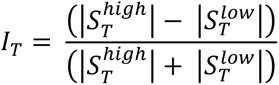

*I*_T_ values ranged between -1 and +1, where increasingly negative and increasingly positive values indicate greater relative sensitivity to torques in the low and high ranges, respectively, and zero indicated that an afferent was equally sensitive in the low and high torque range. A preferred direction in which a given afferent showed maximum torque sensitivity was used for *I*_T_ calculations.

D-prime (d’) analyses were calculated for torque magnitude as:

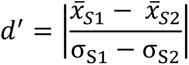

Where x̄ is the mean and *σ* the standard deviation of the number of spikes recorded from an afferent in response to six torque trials of magnitude S1 and S2. For human torque magnitude d’ analyses, s1 was 7.5 mNm and S2 was 3.5 mNm, and for monkey trials S1 was 5.5 mNm and S2 was 2 mNm. For human and monkey torque direction d’ analyses, S1 is anticlockwise torque trials and S2 is clockwise torque trials.

Unless otherwise stated, statistical analyses were performed in R (version 4.0.2; R Core Team, 2020). To assess differences in torque sensitivity and discriminability we used one-way or two-way analysis of variance (ANOVA) or repeated measures ANOVA, where necessary (*anova_test* function, *rstatix* package version 0.7.0; Kassambara, 2021). For one-way non-parametric comparisons the Kruskal-Wallis rank sum test was used *(kruskal_test* function, *stats* package; R Core Team, 2020). Where appropriate, skewed data were log transformed for ANOVA tests. Effect sizes were reported where appropriate, as eta squared (ŋ^2^), a measure of the proportion of variance accounted for by a variable, or F-values. Post hoc pairwise comparisons were performed with the Tukey adjustments for ANOVAs and Bonferroni correction method to adjust for repeated measures ANOVAs. Results are written as mean ± standard error of the mean (SEM), or median and range. The probability selected as significant was *p* < 0.05, for all tests.

## Results

Single-unit recordings were obtained from 39 FA-I, 31 SA-I and 13 SA-II afferents located in the finger pads. First, we analysed the mean and peak discharge rates of FA-I, SA-I, and SA-II afferent responses and first spike latencies in FA-I and SA-I afferents to determine how they were influenced by the magnitude and direction of torque. Next, we analysed how the magnitude of background normal force influenced those afferent responses to torque. Finally, we addressed the question of how afferent receptive field centre (RFC) locations affect their torque responses. As noted above, there have been no previous characterisations of human SA-II afferent responses to torque, so we pay particular attention to characterising SA-II torque response features over the other afferent response properties for which data from monkeys exist (Birznieks et al., 2010; Redmond et al., 2010a; Redmond et al., 2010b; Fu et al., 2012; Khamis et al., 2015).

### SA-II afferent responses to torque

#### Typical SA-II afferent torque response characteristics

SA-II afferents had a variety of response characteristics to different torque magnitudes and directions, and these were also influenced by the background normal force. SA-II afferents have large receptive fields that sometimes encompassed the whole fingertip, and some afferents had an ongoing discharge without the presence of a stimulus (Knibestöl and Vallbo, 1970; Knibestöl, 1975). Typically, SA-II afferents showed increased firing rates at the onset of normal force application, which adapted to a steady level during the normal force plateau. Examples of several distinct SA-II response profiles to torque magnitudes applied in both directions are shown in **Figure 2**. Torque application in one of the two directions often caused SA-II afferents to show pronounced discharge rate increases and their responses could be inhibited by torques applied in the opposite direction. In some SA-II afferents instantaneous discharge rate peaked during the loading phase, after which discharge rate slightly decreased to a steady level during the plateau phase, followed by a change in discharge rate with the opposite sign during the unloading phase (**Figure 2A**). **Figure 2A and B** show afferents whose discharge rates were excited by anticlockwise torques and inhibited by clockwise torques during the loading and plateau phases. **Figure 2C** shows a less common example of an afferent that had an inhibited discharge rate in response to torque in one direction and was not significantly influenced by torque in the opposite direction. In summary, SA-II response behaviours included being (i) excited or inhibited by one torque direction, (ii) excited by one torque direction and inhibited by the opposite direction, or (iii) excited or inhibited by both torque directions.

**Figure 2.**
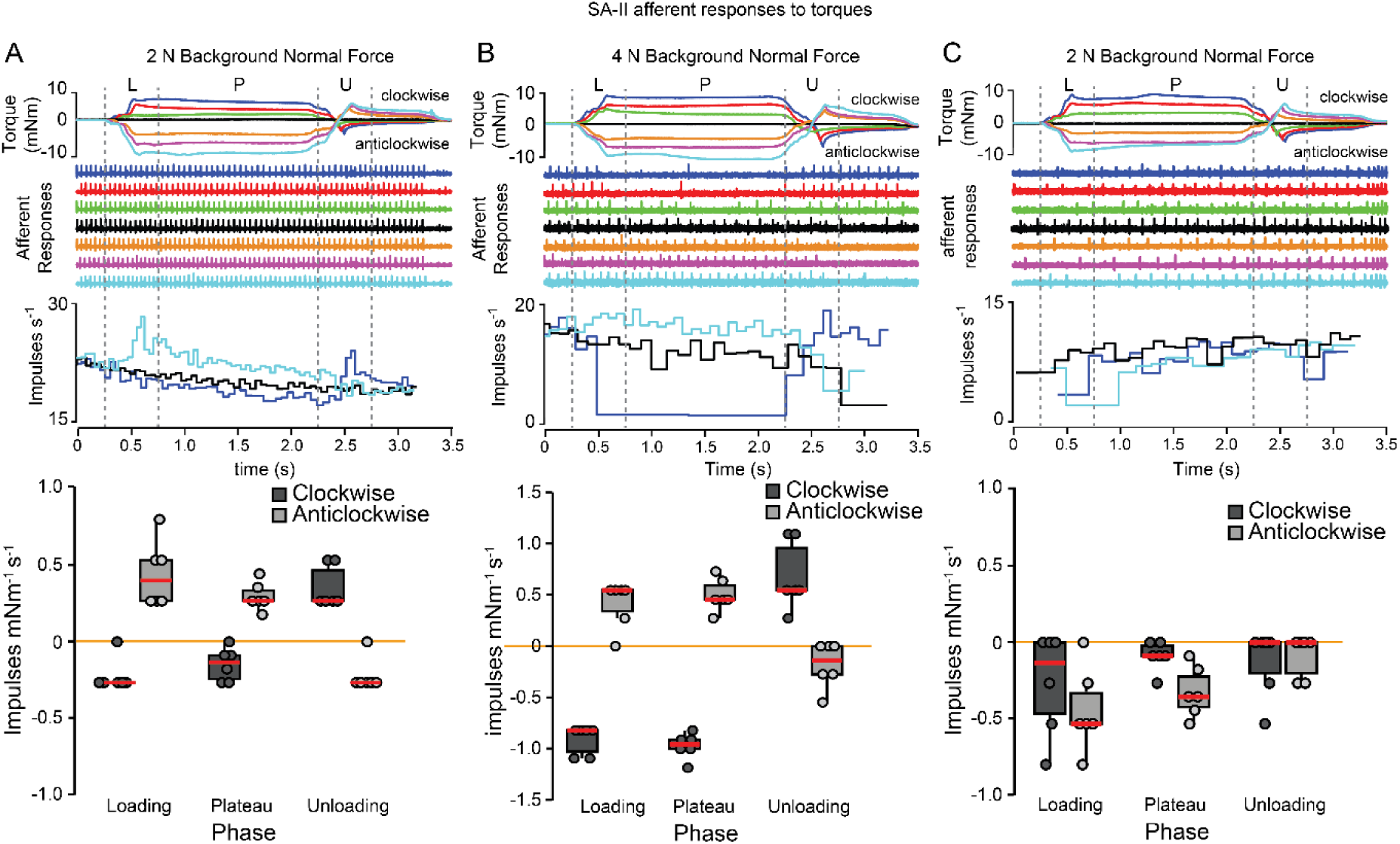
Responses of individual SA-II afferents to torque with background normal force. **A-C**) Example stimuli and responses from three individual SA-II afferents with response profiles showing enhanced discharge rates in response to one of torque directions and suppressed discharge rates to the opposite torque direction (**A-B**). The afferent in **C** showed suppressed discharge rates in response to both torque directions (**C**). The top traces show seven example stimuli applied to the same afferent with torque 3.5 mNm, 5.5 mNm, and 7.5 mNm in clockwise and anticlockwise directions and to normal force only (black trace). Each torque stimulus is an example from a single trial. Below, corresponding spiking activity from single afferent responses to each torque stimulus are colour-coded to match torque stimuli. Each vertical line indicates an afferent action potential (spike). The instantaneous discharge rate in response to 7.5 mNm clockwise and anticlockwise torques, and with normal force only, are shown below the afferent responses. Vertical grey dashed lines separate the torque phases: loading (L), plateau (P), and unloading (U). The box and whisker plots below show the torque sensitivity for opposing torque directions, during each torque phase. Red horizontal lines are medians and box upper and lower bounds show the upper and lower quartiles. Upper whisker shows the largest observation less than or equal to the upper box hinge + (1.5 × inter-quartile range), and the lower whisker shows the smallest observation greater than or equal to the lower box hinge - (1.5 × inter-quartile range). Each filled circle is an individual torque sensitivity (S_T_^all^) value, calculated based on 7.5 mNm torque stimuli. Stimuli were applied six to twelve times, so each plot has a corresponding number of sensitivity values.

Afferent responses that were inhibited during the torque loading and plateau phases were typically excited during the unloading phase. At the end of the unloading phase the stimulation surface was returned to its starting rotational position from which torque was applied. However, this did not result in torque returning to zero—there was always a torque overshoot in the opposite direction, which decreased over time while the surface remained fixed in position indicating that, at least in part, this effect was caused by viscoelastic properties of the skin tissue. It is also possible that, during torque loading, localized ‘slip-and-stick’ events occurred at the skin-contact area edge, particularly at higher torque magnitudes. When the surface was returned to its starting position, these skin regions, which due to ‘slip-and-stick’ events didn’t move all the way with the probe, were now moved back by the full angular distance, thereby pushing those skin regions in the opposite direction, and generating a torque overshoot at the end of torque loading. The torque overshoot in the opposing direction could cause the discharge rate to peak (**Figure 2A, B**). Similarly, afferents with discharge rates excited by torque during the loading and plateau phases often returned to the baseline normal force response rate and might show inhibitory negative peaks during the torque unloading overshoot, although this effect was less frequent. This mechanical phenomenon that causes a hysteresis effect and enhances afferent responses during unloading, seems to be representative of natural movements during object manipulation. In such cases, torque increases and decreases during consecutive stages of manipulation when the object is tilted and then returned to its original orientation.

#### Size of SA-II afferent population influenced by torque

##### Mean discharge rate

Using the mean discharge rate during each torque phase as the response measure we used Spearman’s Rank correlations to determine whether torque magnitude significantly influenced the mean discharge rate. Most SA-II afferents were significantly influenced by torque with 2 N background normal force and had directional differences. For all mean and peak discharge rate comparisons, afferents that were only tested with one torque direction were excluded, so that the proportions of afferents with responses scaled by clockwise and anticlockwise torques could be compared. Moreover, only afferent responses to torques applied with 2 N background normal force were included, as most afferents were stimulated with this background normal force, but not necessarily with higher background normal forces.

*The loading phase.* During the loading phase, 46% (6/13) of SA-II afferents had responses scaled by clockwise torque (five excited, one inhibited), and 69% (9/13) had responses scaled by anticlockwise torque (six excited, three inhibited). Four SA-II afferents had responses scaled in both torque directions, of which two had firing rates that were excited in response to one torque direction and inhibited by the opposite direction, and two were excited by both torque directions. Altogether, 85% (11/13) of SA-II afferents were scaled by torque magnitude in at least one direction during the loading phase (**Figure 3A**). In some trials, afferent responses with torque were significantly different only from trials with normal force but did not show discernible response scaling to the torque magnitudes in each direction, within the magnitude range tested in this study. We refer to such afferents as having a *non-graded response to torque* in each direction, which were determined using the Mann-Whitney- Wilcoxon test. In total, 100% (13/13) SA-II afferents were either scaled or had non-graded responses to torque in at least one direction. Three afferents that had torque-scaled excitatory responses to one torque direction had inhibitory non-graded responses to the opposite direction (*p* < 0.05, Mann-Whitney-Wilcoxon test). Including graded and non-graded responses, SA-II afferents had strong torque direction selectivity among the population, as five afferents’ excitatory-inhibitory response switched between opposite directions, seven responded with excitation or inhibition to torque selectively in one of the directions, and only two out of thirteen seemed to be indifferent and were excited by torque in either direction.

**Figure 3.**
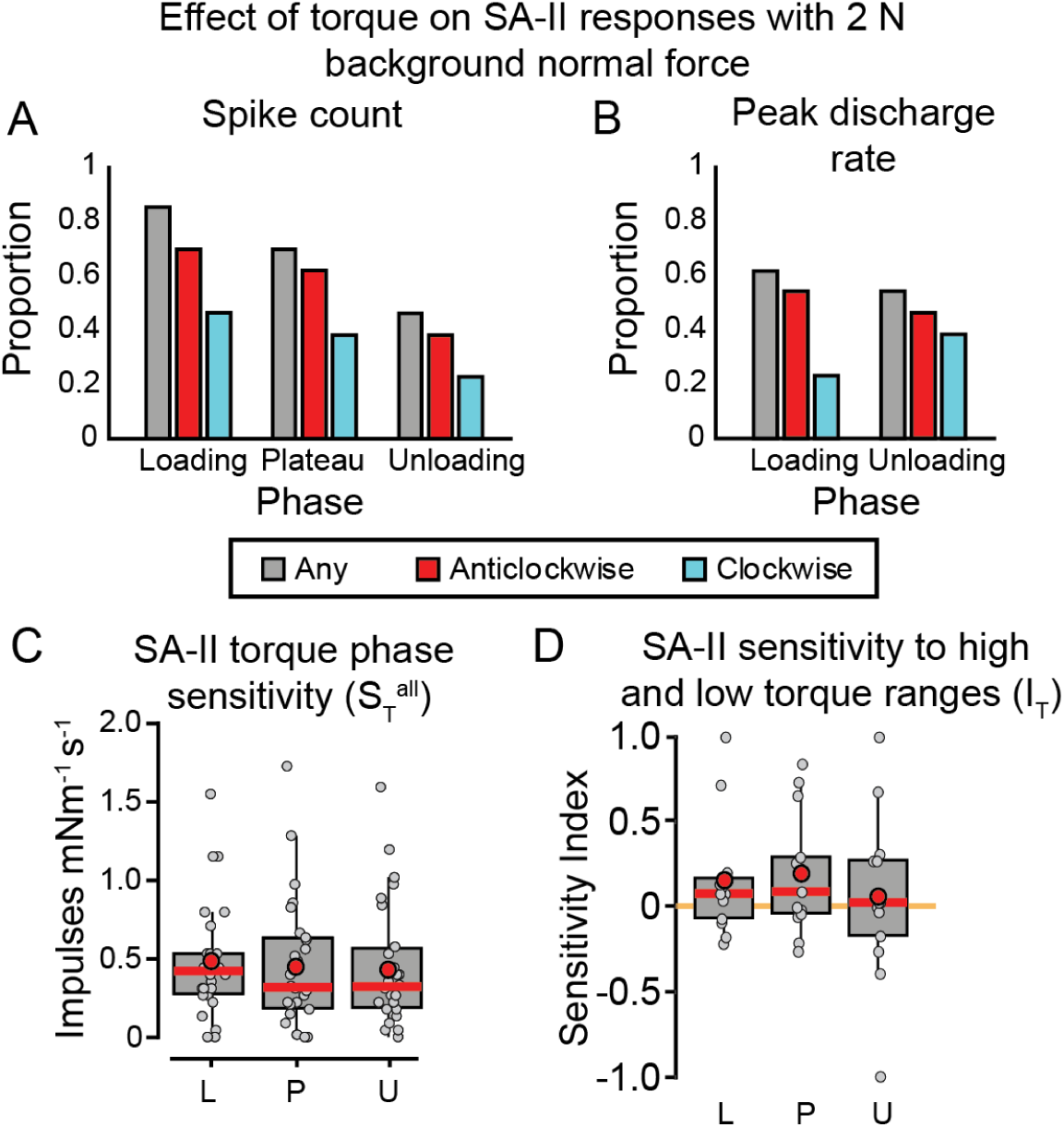
Effect of torque on SA-II afferents in each torque phase. **A-B**) The proportion of SA-II afferents with scaled (**A**) mean or (**B**) peak discharge rates in response to torque, with 2 N background normal force. Spike counts (**A**) were quantified for each torque phase, but peak discharge rates (**B**) were only quantified for the dynamic loading and unloading phases. **C**) The torque sensitivity (*S*_T_^all^) of each afferent in each torque phase shown in box and whisker plots. Red circles and horizontal lines are means and medians, respectively. Each filled circle is the mean S_T_^all^ for individual afferents. **D**) The torque sensitivity index *I*_T_ estimated at 2 N background normal force comparing S_T_^low^ and S_T_^high^ representing afferent response changes within the low torque range (0- 3.5 mNm) and the high torque range (3.5-7.5 mNm), respectively, is shown in box and whisker plots (see Figure 2 for details). Positive values indicate the afferent responses were more strongly modulated within the high torque range and negative values indicate the afferents were more strongly modulated within the low torque range. Each afferent scaled by torque is represented by one sensitivity index *I*_T_ in the direction which gave the larger absolute value. Red horizontal lines and circles indicate medians and means, respectively. Orange horizontal line indicates zero for comparison.

*The plateau phase.* Torque scaling was similar during the plateau phase, as 38% (5/13) of SA- II afferents had mean discharge rates scaled by clockwise torque magnitude (three excited, two inhibited), and 62% (8/13) were scaled by anticlockwise torques (six excited, two inhibited). Only 5/13 (39%) SA-II afferent responses were scaled in both torque directions, of which two had excitatory-inhibitory responses switch between opposite directions, one was excited and one inhibited in both torque directions. Altogether, 85% (11/13) of SA-II afferents were scaled in at least one direction during the plateau phase (**Figure 3A**). When non-graded responses to torque were taken into account, then all (13/13) SA-II afferents were either scaled or had non-graded responses to torque in at least one direction.

*The unloading phase*. Fewer SA-II afferents had responses scaled by torque during the unloading phase, with 23% (3/13), and 38% (5/13) of afferents scaled by clockwise or anticlockwise torque, respectively, and 46% (6/13) scaled in at least one direction. When non-graded responses to torque were considered, 12/13 SA-II afferents were either scaled or had non-graded response to torque in at least one unloading direction.

#### Peak discharge rate

We also determined the proportion of afferents in which peak discharge rates were influenced by torques during the loading and unloading phases. If discharge rates were inhibited, we measured the minimum discharge rate at the trough. The effect of torque on SA-II afferents’ peak discharge rates was typically similar to the effect on mean discharge rate (**Figure 3A, B**). In the torque loading phase, peak discharge rates were scaled in 23% (3/13) of SA-II afferents in response to clockwise torques during the loading phase, and in 54% (7/13) by anticlockwise torque, of which two afferents had excitatory-inhibitory responses switch between opposite directions. Altogether, 62% (8/13) of SA-II afferents had peak discharge rates scaled by torque magnitude in either direction.

During the unloading phase, clockwise and anticlockwise torques scaled 38% (5/13) and 46% (6/13) of SA-II afferent peak discharge rates, respectively, and altogether 54% (7/13) were scaled by torque in either direction.

#### SA-II torque sensitivity comparison

Next, we sought to determine whether there was a preference for how afferents respond to stimuli during dynamic (loading and unloading) phases, and during the static plateau phase. The torque sensitivity index, *S*_T_^all^, expressed as impulses per mNm per second, was calculated as changes in mean discharge rate between no torque and the largest torque magnitude tested (**Figure 3C**). We found no significant differences in torque sensitivity among the three stimulation phases for the population of SA-II afferents (*p* = 0.71, *F*(2,180) = 0.34, repeated measures two-way ANOVA). It is possible that asymmetries in the fingertip structure could influence torque direction sensitivity. We compared SA-II sensitivity to torques in opposite directions and found that there was no significant differences (*p* = 0.45, *F*(1,180) = 0.4, repeated measures two-way ANOVA).

In the following analyses we wanted to find out which is the more optimal torque range tested in the current study. Some afferents might be less sensitive to low torques and respond preferentially to higher torques, others might saturate responses between two higher torque levels thus having better discrimination at a lower torque range. To determine whether the stimulus-response curve is expected to show accelerating increasing sensitivity or decelerating and showing signs of saturation, we calculated the torque sensitivity increase index, *I*_T_, based on torque sensitivity indices estimated for low *S*_T_^low^ (0-3.5 mNm) and high *S*_T_^high^ (3.5-7.5 mNm) torque ranges (**Figure 3D)**. Positive values indicate the afferents were increasingly more sensitive to torque differences in the high torque range, while negative values indicate the afferents were more sensitive to torque differences in the low torque range and saturating responses at higher torque levels. Individual SA-II afferents in a population had highly variable sensitivities to change in the high and low torque ranges, with the median values typically close to zero during the loading and plateau phases, suggesting they had similar sensitivity to high and low torques (**Figure 3D**). We found there was significant differences in *I*_T_ among torque phases (*p* = 0.01, *F*(2,94) = 4.7, one-way repeated measures, ANOVA) (**Figure 3D**), but post-hoc comparisons revealed that the difference was only between the loading (0.15 ± 0.09) and unloading (0.06 ± 0.13) phases (*p* = 0.03 Bonferroni adjusted).

### Effect of normal force on SA-II torque responses

#### Individual SA-II afferent responses

The torque sensitivities, S_T_^all^, of many individual SA-II afferents were clearly influenced by the background normal force. An example afferent had high torque sensitivity with 2 N background normal force (**Figure 4A**), but the torque responses were greatly reduced with higher background normal force 3 N (**Figure 4B**) and 4 N (**Figure 4C**). The effect of the background normal force was significant (*p* < 0.001, F(2,108) = 21.8, three-way repeated measures ANOVA). Another interesting example of an SA-II afferent with responses that were influenced by background normal force is shown in **Figure 4D-F**. With 2 N background normal force, this afferent showed increased discharge rate during the loading and plateau phases to torques in both directions but, when the background normal force was increased to 3 N (**Figure 4E**) and 4 N (**Figure 4F**), the effect of anticlockwise torques switched from excitation to inhibition. The effect of force on torque sensitivity S_T_^all^ was significant in this afferent (*p* < 0.001, F(2,90)= 13.4, three-way repeated measures ANOVA). The lowest sensitivity in response to torque was in the presence of 3 N in comparison to 2 N and 4 N background normal force as it was the midpoint at which the transition from excitatory to inhibitory pattern occurred. These examples suggest that background normal force is important in determining the torque sensitivity of an afferent and increasing the normal force can either increase or decrease torque sensitivity of SA-II afferents. It is likely that fingertip geometry and mechanics, in combination with the receptive field location determines those complex interactions between the afferent torque and normal force responses. It must be noted that SA-II afferents have relatively large receptive fields, and the location of the receptor ending often cannot be precisely determined. For these two example afferents the receptive fields were relatively large, covering a large part of the fingertip with diffuse borders that were hard to demarcate; however, we were able to determine the location of the receptor endings: for the afferent in **Figure 4A-C** it was located 10 mm distal to the stimulus rotational centre at the fingernail border, whereas for the afferent shown in **Figure 4D-F** it was located 10 mm proximal near the midline from the stimulus rotational centre.

**Figure 4.**
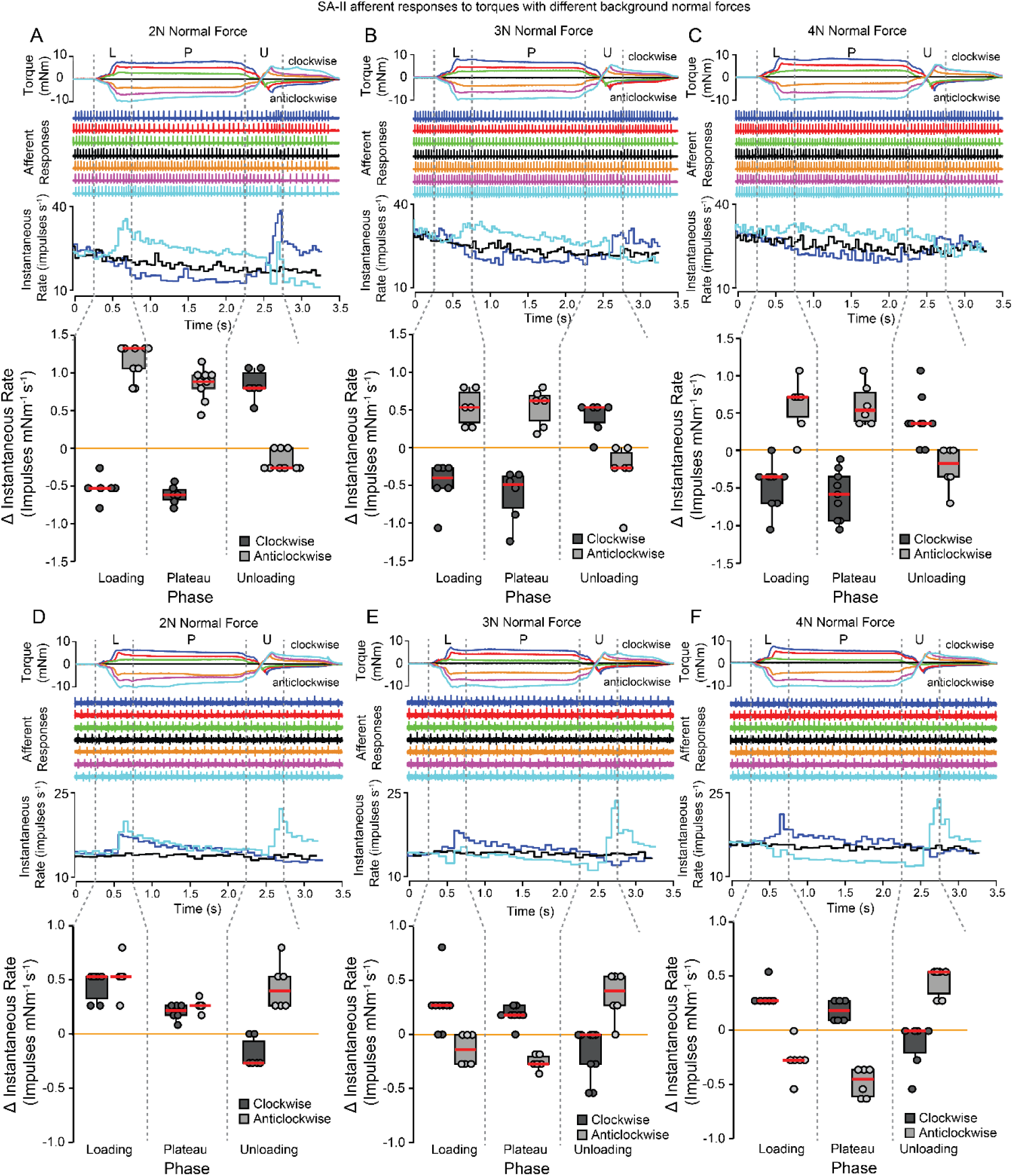
The effect of background normal forces on SA-II afferent responses to torque. Responses to torque from the same single afferent are shown when the torques were applied with 2 N (**A**), 3 N (**B**), and 4 N (**C**) background normal forces. Note that in this afferent the torque sensitivity appears to decrease with increasing background normal force. **D-F)** Torque responses from another afferent are shown with three different background normal forces. Note that this afferent’s sensitivity to clockwise torques appears to remain the same with increasing background normal forces, but in response to anticlockwise torques when background normal force increased from 2 N (**D**) to 4 N (**F**), torque effects transitioned from excitation to inhibition. See Fig 2 legend for detailed description.

#### SA-II afferent population effects

The responses of exemplified single afferents shown in **Figure 4** indicate that background normal force may substantially influence SA-II afferent sensitivity to torque. We sought to investigate what effect the background normal force had across the SA-II afferent population. There was a clear relationship showing that the proportion of afferents with responses scaled by torque during the loading phase was inversely related to background normal force (**Figure 5A**). During the plateau and unloading phases, the relationship was less clear, but in both cases, the proportion of afferents with responses scaled by torque were greater with 2 N background normal force than with 4 N, as was the case during the loading and unloading phase when we used peak discharge rate as the response measure (**Figure 5B**).

**Figure 5.**
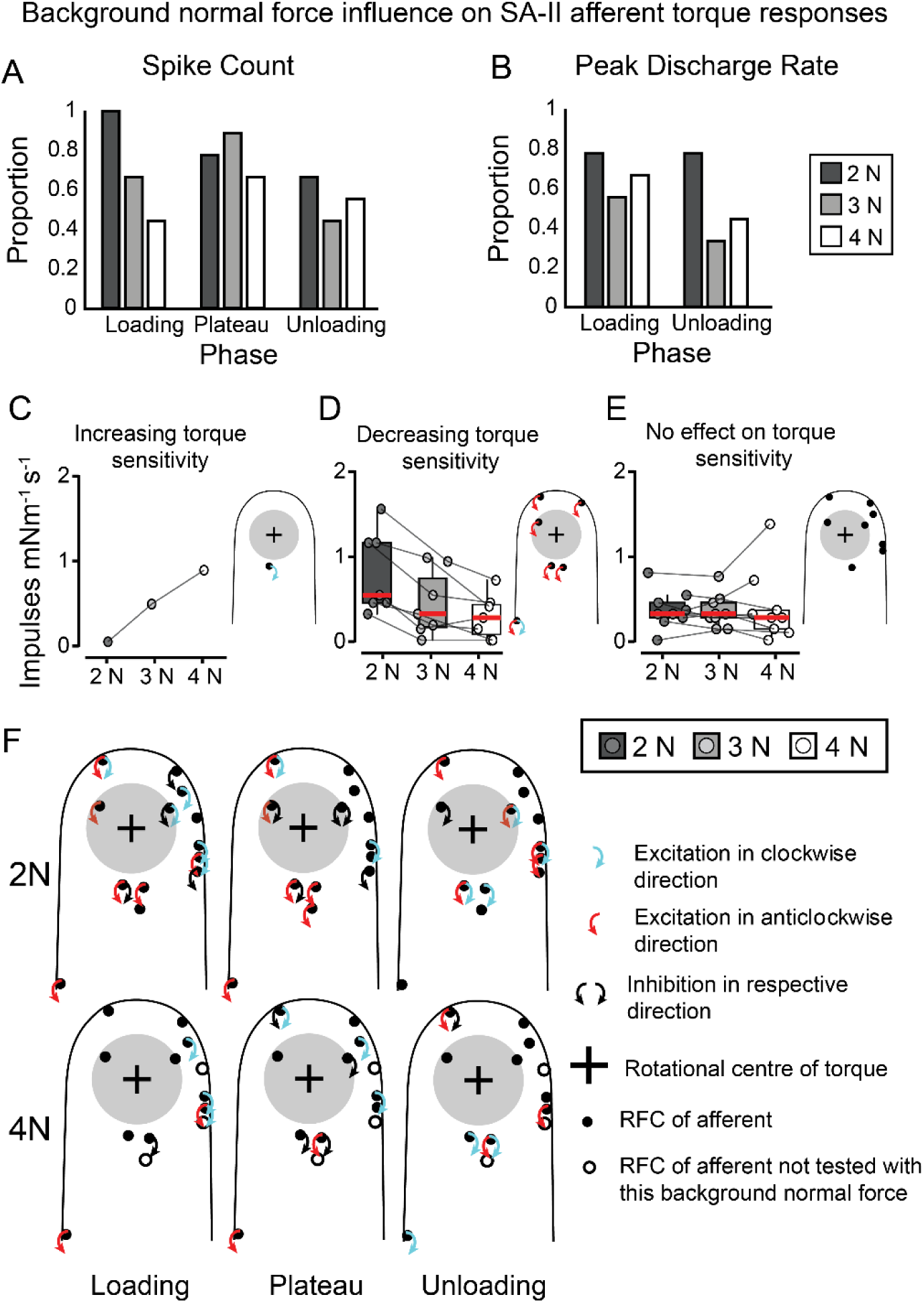
Effect of normal force and receptive field centre location on SA-II afferent’s torque responses. **A-B**) The proportion of SA-II afferents that showed responses that were scaled by torque in at least one direction as measured by (**A**) spike count and (**B**) peak discharge rate, for each normal force. Spike counts (**A**) were quantified for each torque phase, but peak discharge rates (**B**) were only quantified for the dynamic loading and unloading phases. **C-E**) SA-II torque sensitivities with 2, 3, and 4 N background normal forces, separated by afferents that had significantly (**C**) increasing or (**D**) decreasing torque sensitivity with increasing background normal force, or were unaffected by background normal force (**E**). Only afferents that were tested with torque at each of the three normal forces are included. Values from individual afferents are connected by lines. The fingertip contours to the right of panels indicate the receptive field centre (RFC) of these afferents. Arrows show the direction of sensitivity to torque between 0-7.5 mNm (S_T_^all^) in the torque loading phase. **F**) The RFC location of each afferent and the direction in which its response was scaled by torque are plotted for each torque phase across a standard fingertip. The top row shows the spatial relationship of SA-II torque scaling, with 2 N background normal force, and the bottom row with 4 N background normal force.

To determine whether individual afferent sensitivity measures to torque were scaled by the normal force on which torque stimuli were superimposed, we used Spearman’s Rank correlation analyses. Afferents that were tested with all three normal forces (n = 10) were used in these analyses. We investigated this relationship only during the torque loading phase, as this is when encoding torque changes is most important for object manipulation and lifting. In **Figure 5C-E** we separated the afferents into three groups; in which normal force increased, decreased, or had no effect on torque sensitivity. We found that 60% (6/10) of SA- II afferents were significantly scaled by background normal force in at least one direction (median correlation coefficient = -0.37, range = -0.79 – 0.77), of which all six had decreasing torque sensitivity with increasing background normal force, but one of these afferents also had increasing torque sensitivity for torques in the opposite direction. The distribution of SA- II afferent RFCs over the distal fingertip is illustrated for groups of afferents with increasing, decreasing, or no effect on torque sensitivity (see schematic fingertips to the right of **Figure 5C-E**).

#### Relationship between receptive field location and SA-II torque sensitivity

The SA-II afferents had approximate RFCs across most of the distal phalanx and only one of these afferents’ responses was not influenced by torque, suggesting that, regardless of the RFC location on the fingertip, SA-II afferents were capable of signalling changes in torque (**Figure 5F**). Torsional loads exerted on the skin increase with distance from the stimulus rotational centre up to, and potentially just beyond, the edge of the contact area. Consequently, cutaneous mechanoreceptors may experience different mechanical effects at different distances from the rotational centre. Near the centre, the skin is more likely to be in safe contact with the surface, so the skin is moved relative to the deeper tissue. Further, at the edge of the stimulus contact there is a higher probability that the skin is in contact but is subjected to ‘slip and stick’ events. Just outside the contact area the skin is exposed to significant shear forces as the skin within the contact area is moved relative to the skin outside the contact area. However, the exact distances where these phenomena occur would be difficult to determine and would depend on multiple factors outside experimental control, for example, skin compliance and variation in friction between the skin and surface. In general, it could be expected that *S*_T_^all^ would increase with RFC distance from the stimulus centre, up to a point somewhere just outside the contact area where S_T_^all^ would likely decline. Visual inspection of data did not reveal any such trend as afferents in any location on the fingertip were sensitive to torque possibly due to the large size of SA-II afferent receptive fields. Further quantitative assessment of this effect was outside the scope of this study as it would require a dedicated experimental setup and larger samples of afferents to perform biomechanical analyses of fingertip deformation, for example, using fingerprint image processing

### SA-I afferent responses to torque

#### Typical SA-I afferent torque response characteristics

SA-I afferents had small and discrete receptive fields, and were excited by normal force application, showing a rapidly increasing discharge rate at the onset, which either continually declined or declined to a steady state during the normal force plateau. At torque onset, SA-I afferents most frequently showed excitatory response, but we also observed some SA-I afferents in which torque inhibited their response to background normal force. **Figure 6** shows example responses of a typical SA-I afferent to three torque magnitudes applied in the clockwise and anticlockwise directions, superimposed on 2 N normal force. During the loading and unloading phases, this afferent was excited by both clockwise and anticlockwise torques. During the plateau phase, only the clockwise torques caused clear enhancement of the discharge rate (**Figure 6**).

**Figure 6.**
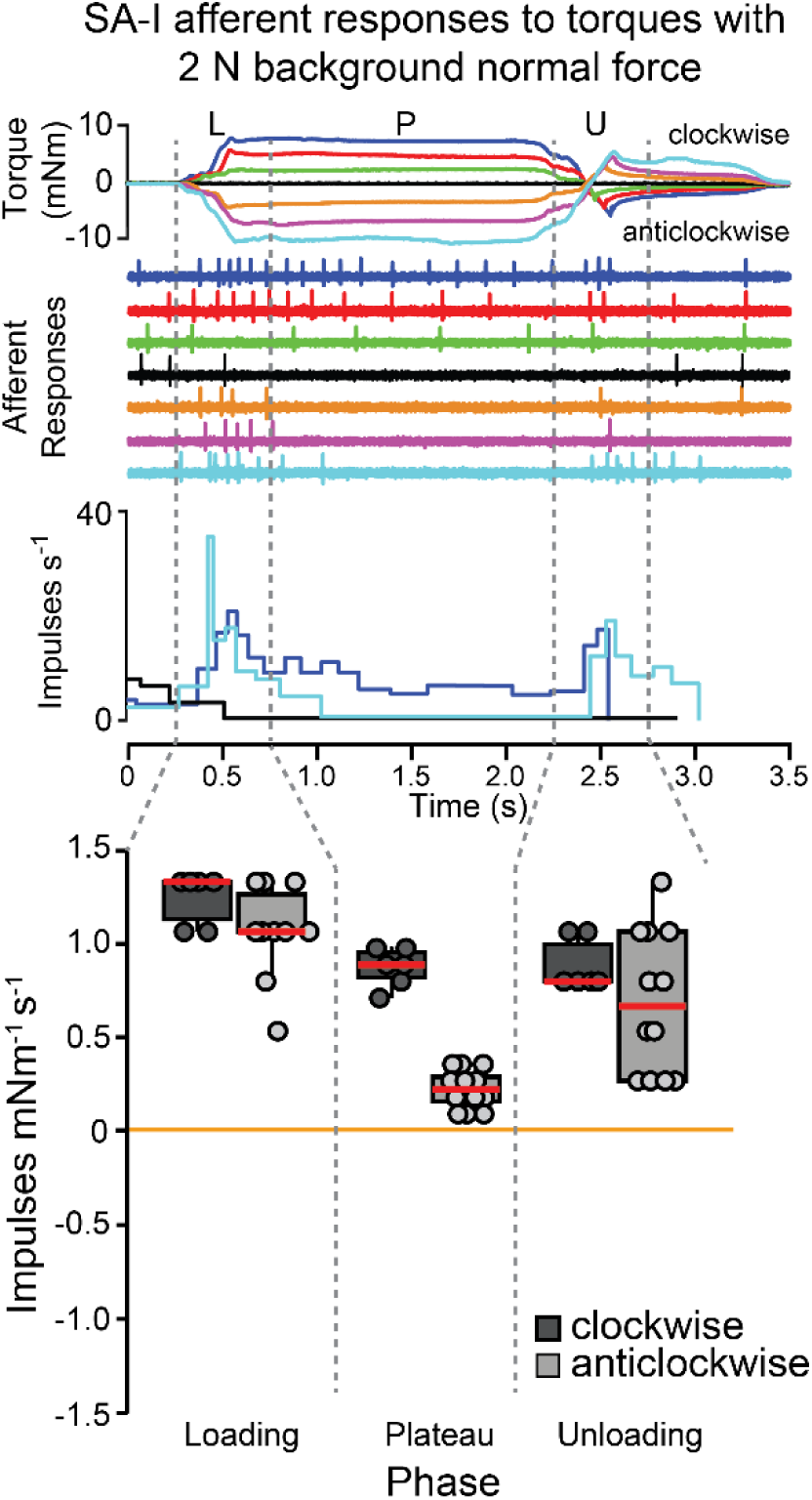
Example of a single SA-I afferent response to torques with 2 N background normal force. The SA-I afferent’s responses show dynamic enhanced responses in the loading and unloading phases to torques applied in both directions. In the plateau phase, the afferent’s responses show an enhanced discharge rate to clockwise torques and a weakly enhanced response to anticlockwise torques. See Fig 2 legend for detailed description.

#### Size of SA-I afferent population influenced by torque

##### Mean discharge rate

In most SA-I afferents mean discharge rate was scaled by torque superimposed on 2N normal force. During the loading phase, 45% (14/31) of SA-I afferents’ responses were scaled by clockwise torques, of which none were inhibited, and 58% (18/31) had responses scaled by anticlockwise torques, of which two were inhibited (**Figure 7A**). Altogether, 65% (20/31) of SA-I afferents had mean discharge rates scaled by torque in at least one direction during the loading phase from which 12 afferents’ responses were scaled by both torque directions (**Figure 7A**). Two of the non-scaled afferents had non-graded responses to torque.

**Figure 7.**
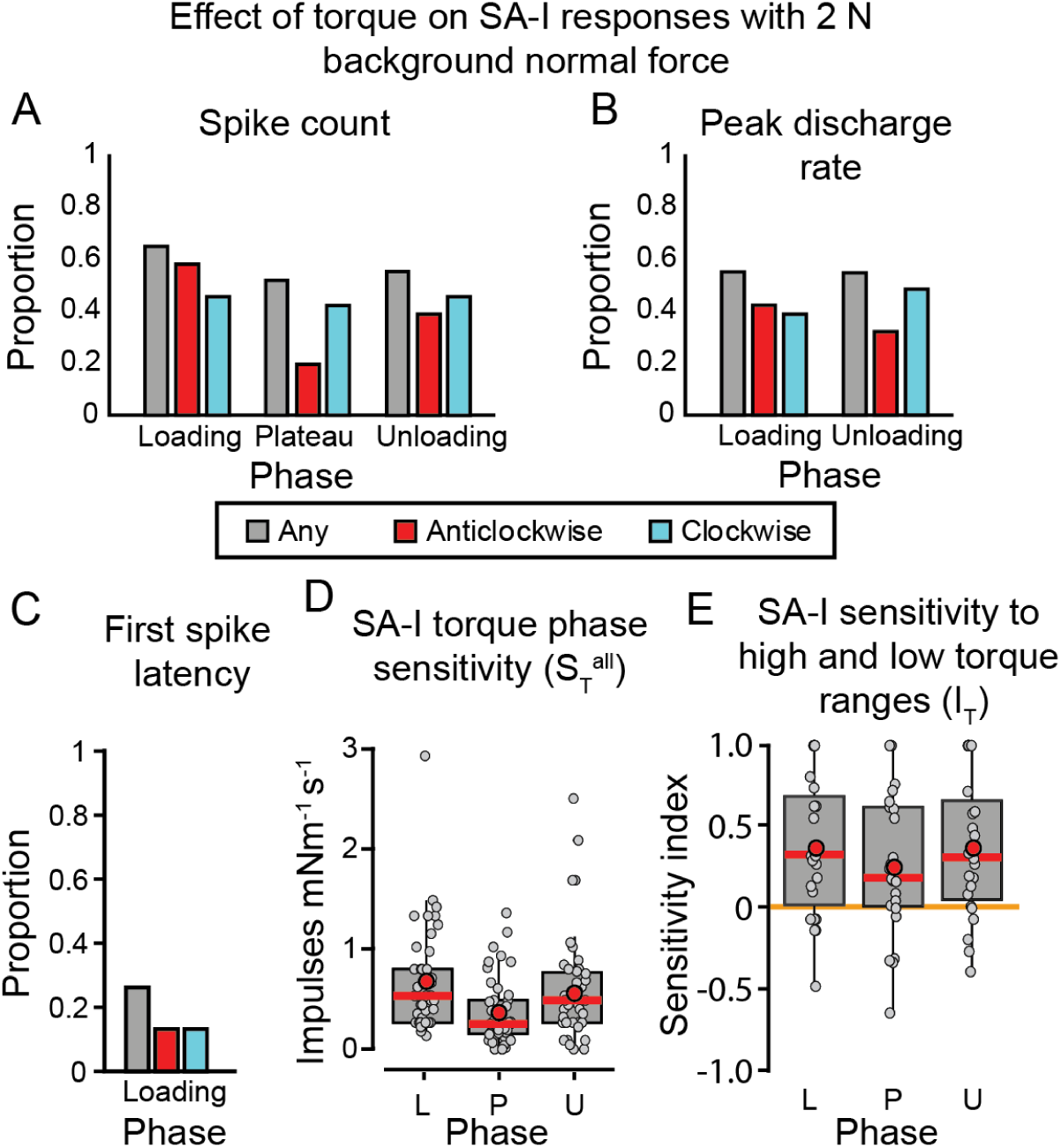
Effect of torque on SA-I afferents in each torque phase. **A-C)** The proportion of SA-I afferents that showed responses that were scaled by torque in at least one direction as measured by (**A**) spike count, (**B**) peak discharge rate, and (**C**) first spike latency, with 2 N background normal force. **D)** The torque sensitivity (*S*_T_^all^) of each afferent in each torque phase, with 2 N background normal force is shown in box and whisker plots. **E**) The torque sensitivity index comparing S ^low^ and S ^high^ for each afferent in each torque phase, with 2 N background normal force is shown in box and whisker plots. See Figure 3 legend for detailed description.

Proportions of SA-I afferents with torque scaled responses were slightly reduced during the plateau phase, as 42% (13/31) of afferents had responses scaled by clockwise torque, of which three were inhibited, and 19% (6/31) by anticlockwise torques, of which three were inhibited (**Figure 7A**). Altogether, 52% (16/31) of SA-I afferents had responses that were scaled by torque in at least one direction during the plateau phase from which only 3 were scaled in both directions (**Figure 7A**) and six scaled only in one torque direction had non-graded responses to the opposite torque direction. In two afferents only non-graded responses to torque in one of directions could be detected.

During the unloading phase, 55% (17/31) of SA-I afferents had responses that were scaled by torque in at least one direction during the unloading phase, (14 clockwise; 12 anticlockwise and 9 in both directions; **Figure 7A)**. Four of the non-scaled afferents had non-graded responses to torque (*p* < 0.05, Mann-Whitney test).

##### Peak discharge rate

Torque influences on SA-I afferent peak discharge rates were similar to mean discharge rates (**Figure 7A, B**). During the loading phase altogether 55% (17/31) of afferents’ responses were scaled by torque in at least one direction (12 clockwise, 13 anticlockwise and 8 in both directions; **Figure 7B**). The response in the same number of afferents 55% (17/31) was scaled by torque in at least one direction during the unloading phase (**Figure 7B**).

##### First spike latency

Encoding mechanisms such as the recruitment order of tactile afferents might be a fast way to signal torque parameters at the fingertips. To investigate whether first spike latencies depend on torque, we calculated Spearman’s Rank correlations between the latency to the first spike in the torque loading phase and torque magnitude. First spike latencies were only calculated for torques applied with 2 N background normal force. Loading phase first spike latencies were scaled in 13% (4/31) of SA-I afferents, in response to either clockwise or anticlockwise torques, and none of the SA-I afferents’ first spike latencies were scaled by both torque directions (**Figure 7C**). Altogether, 26% (8/31) of SA-I afferents had first spike latencies scaled by torque in at least one of directions (**Figure 7C**). On average, SA-I afferent first spike latencies were 50.3% shorter in response to 7.5 mNm compared to 3.5 mNm torque. This analysis was not performed for the SA-II afferents given their spontaneous discharge.

#### SA-I torque sensitivity comparison

Next, we evaluated SA-I afferent responses to torque by calculating *S*_T_^all^ based on mean discharge rate for each torque phase, direction, and background normal force (**Figure 7D**). SA-I afferents torque sensitivity varied significantly between the torque phases (*p* < 0.001, F(2,318) = 11.8, two-way repeated measures ANOVA) (**Figure 7D**). Pairwise comparisons revealed that SA-I torque sensitivity was significantly greater during the loading and unloading phases compared to the plateau phase (*p* < 0.001, Bonferroni adjusted), but there was no difference between the loading and unloading phases. On average, there were no significant differences in the mean sensitivity of the SA-I afferent population when torques were applied in the clockwise or anticlockwise directions (*p* 0.12, *F*(1,318) = 2.4, two-way repeated measures ANOVA). To determine whether SA-I afferents had different levels of sensitivity with low torque (0-3.5 mNm) and high torque (3.5-7.5 mNm) ranges, we calculated the torque sensitivity increase index (*I*_T_) (**Figure 7E)**. Generally, SA-I afferents had highly variable *I*_T_ values indicating that individual SA-I afferents had preference for, and their highest sensitivity encoding over, the whole range of tested representative torques. There was no difference between phases in this respect (*p* = 0.99, *F*(2,153) = 0.01, one-way repeated measures ANOVA) (**Figure 7E**).

#### Effect of normal force on SA-I afferent torque responses

The background normal force influenced the number of SA-I afferents that had mean discharge rates scaled by torque magnitude. When using mean discharge rate as the response measure, there were fewer afferents scaled by torque when normal force increased during the loading and plateau phases (**Figure 8A**). This was also apparent during the loading phase when peak discharge rates (**Figure 8B**) and first spike latencies (**Figure 8C**) were used as the response measure.

**Figure 8.**
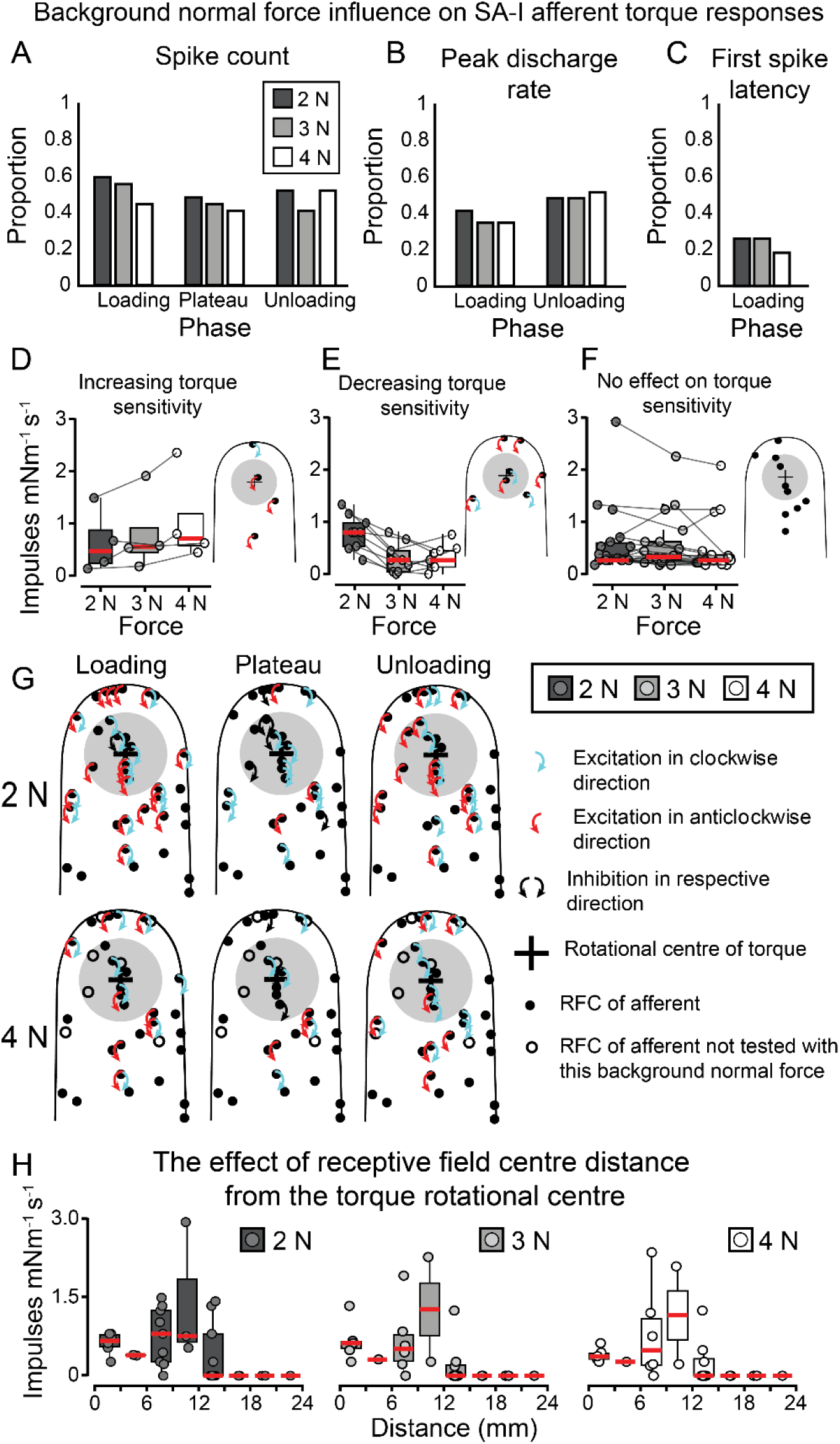
Effect of normal force and receptive field centre location on SA-I afferent’s torque responses. **A-C)** The proportion of afferents that were scaled by torque in at least one direction as measured by (**A**) spike count, (**B**) peak discharge rate, and (**C**) first spike latency for each normal force. **D-F**) SA-I torque sensitivities with 2, 3, and 4 N background normal forces, separated by afferents that had significantly decreasing (**D**) or increasing (**E**) torque sensitivity with increasing background normal force, or were unaffected by background normal force (**F**). **G**) The RFC location of each afferent and the direction of torque scaling are plotted for each torque phase across a standard fingertip. **H**) Sensitivity values from the torque loading phase are shown for SA-I afferents against their RFC distance from the stimulus rotational centre. S_T_^all^ was calculated from the maximum torque sensitivity value between either clockwise or anticlockwise torque directions. Plots are separated by the background normal force. SA-I afferents showed increasing torque sensitivity with distance from the stimulus rotational centre, up to 12 mm, and the most pronounced effect was from 6-12 mm. SA-I afferents with RFC greater than 15 mm from the stimulus rotational centre were unresponsive to torque. See Figure 5 legend for detailed description.

To visualise the effect of force on individual afferents within the population, we separated afferents into three groups; those that showed increasing, decreasing, or no effect on torque sensitivity, respectively (**Figure 8D-F**) with increasing background normal force. Only afferents in which the effect of torque was tested with all three background normal forces were included in this analysis (n = 17). We found that 53% (9/17) of SA-I afferents were significantly scaled by background normal force (median correlation coefficient = -0.21, range = -0.84 – 0.87), of which 78% (7/9) had decreasing torque responses with increasing background normal force, and 44% (4/9) had increasing torque responses with increasing background normal force. Interestingly, three of these SA-I afferents had increasing torque sensitivity in one torque direction and decreasing in the opposite direction, when background normal force increased. There was no difference in the distance of SA-I afferents’ RFC from the rotational centre whether they had increasing (mean distance = 7.0 mm, S.D. = 5.4 mm) or decreasing (mean distance = 7.2 mm, S.D. = 4.5 mm) torque sensitivity with background normal force, and no clear relationship was observed in the distribution of their RFC over the distal part of the fingertip (**Figure 8D and E**).

#### SA-I relationship between receptive field location and torque sensitivity

The RFC of SA-I afferents were located over the entire distal phalanx (**Figure 8G**). Further away from the rotational centre the ratio between the local normal force to local tangential torque decreases thus increasing the likelihood of torsional slips between the skin and the surface. As noted above, the shear forces experienced by the skin are also greatest just outside the edge of the contact area, hence we expected to see torque sensitivity (S_T_^all^) increase with SA- I RFC distances from the stimulus centre. **Figure 8H** shows the torque loading phase sensitivity with three background normal forces. Each SA-I afferent was binned using the RFC distance from the stimulus centre in 3 mm bins and the torque sensitivities plotted for each bin. SA-I afferents had lower torque sensitivity when their RFC was within a 6 mm radius of the stimulus centre, which was the approximate contact area of the stimulus probe, but there appeared to be a greater number of afferents with a high torque sensitivity between 6-12 mm from the stimulus centre, which reduced with higher distances. There were no SA-I afferents sensitive to torque with RFC’s greater than 15 mm from the stimulus centre.

### FA-I afferent responses to torque

#### Typical FA-I afferent torque response characteristics

As expected, FA-I afferents responded to normal force and torque magnitudes exclusively during the dynamic phases of stimulation (loading and unloading phases). An example of a torque response profile from a typical FA-I afferent is shown in **Figure 9**. This afferent’s discharge rate increased by torque magnitude applied in both directions during the loading and unloading phases, but it was silent during the torque plateau phase. As FA-I afferents are not activated by a steady background normal force, they cannot show inhibition by torque.

**Figure 9.**
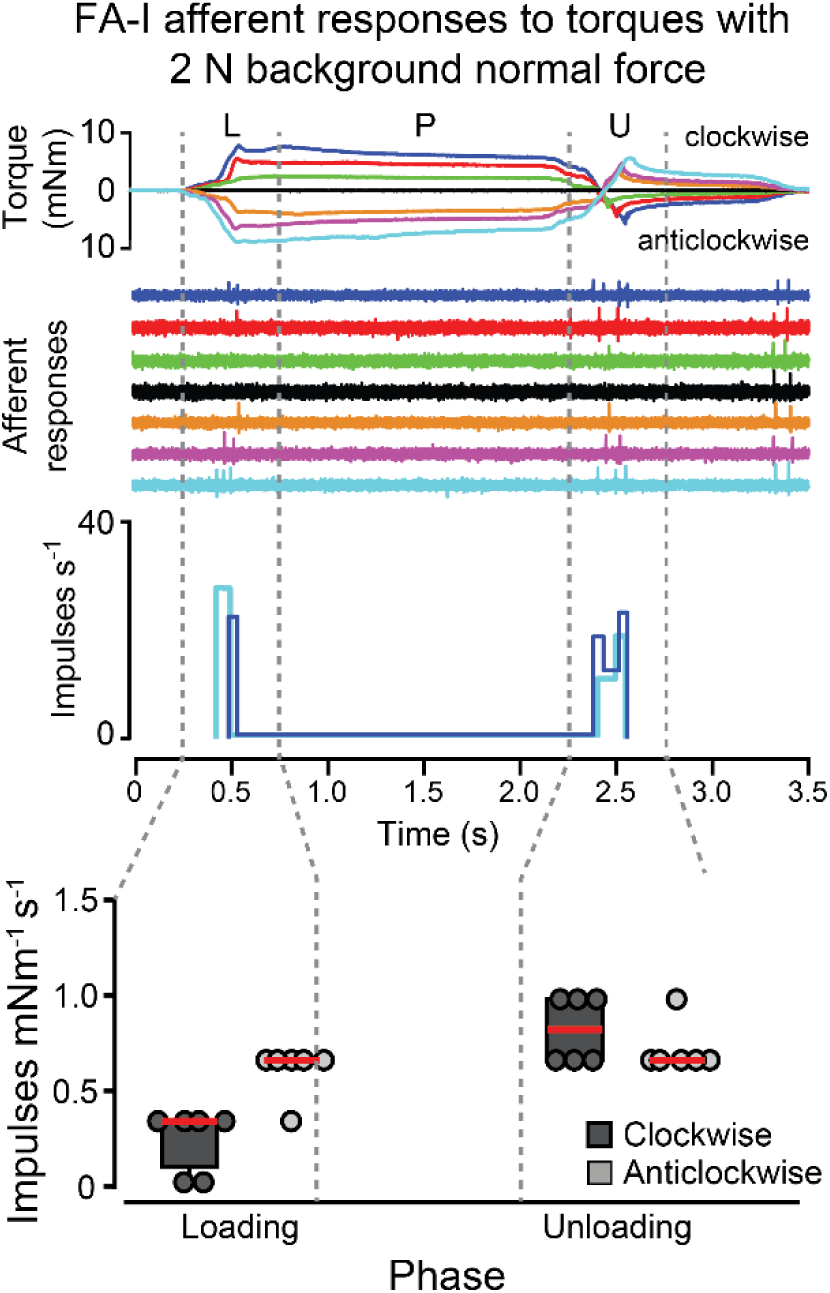
Example of a single FA-I afferent responses to torques with 2 N normal force. The FA-I afferent’s responses show some scaling in the loading and unloading phases to torques applied in both directions. FA-I afferents did not respond in the plateau phase, so this phase has been excluded from analysis. See Figure 2 legend for detailed description.

#### Size of FA-I afferent population influenced by torque

##### Mean discharge rate

In most FA-I afferents the mean discharge rate was scaled by torque during the loading and unloading phases. During the loading phase, 56% (22/39) of FA-I afferents’ responses were scaled by torque (15 in clockwise, 21 in anticlockwise and 14 in both directions; **Figure 10A**). Five FA-I afferents that had torque-scaled responses to one torque direction, had non-graded responses to the opposite torque direction, and one that wasn’t scaled by torque at all, had non-graded responses to anticlockwise torques (*p* < 0.05, Mann-Whitney-Wilcoxon test).

**Figure 10.**
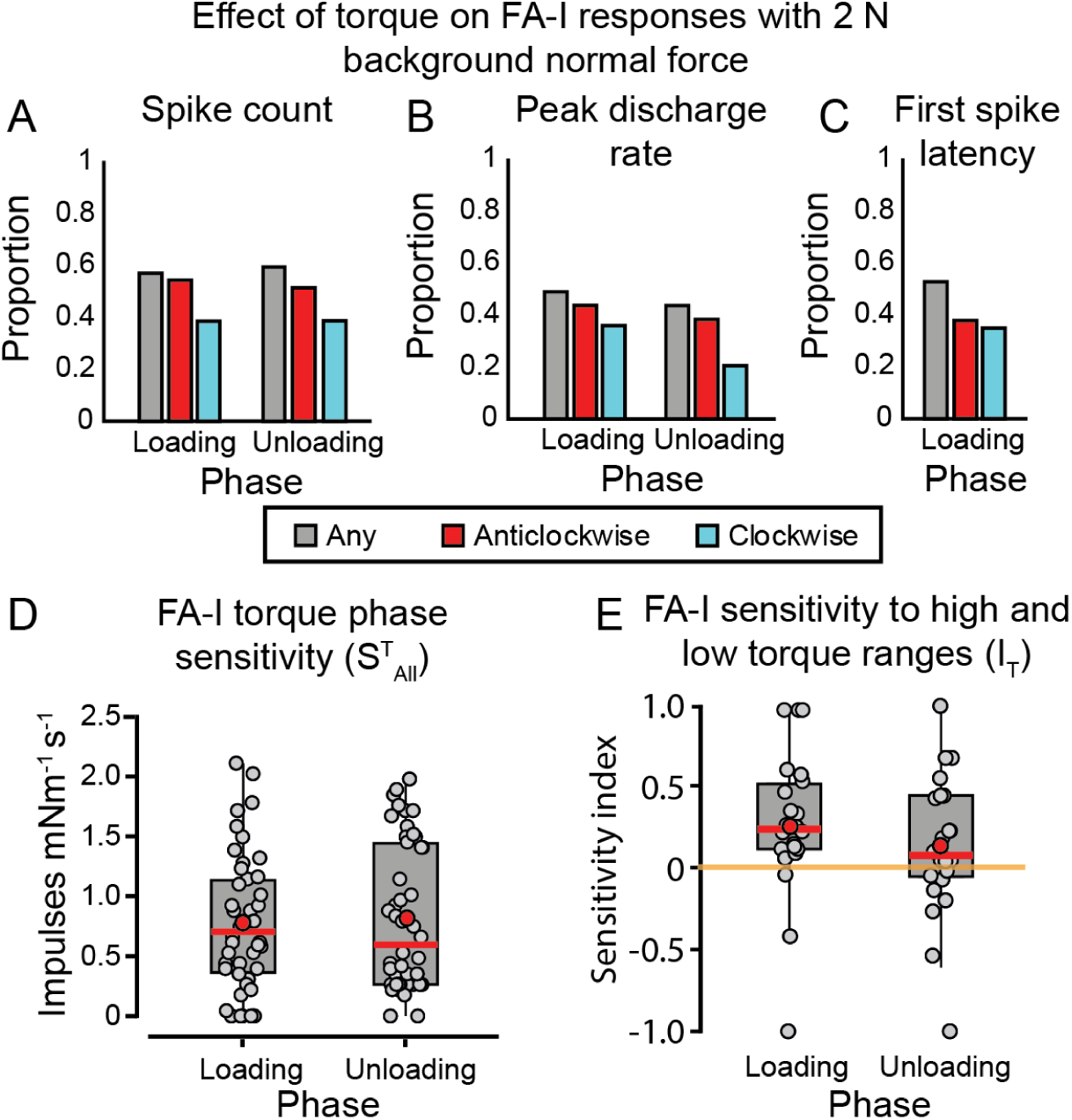
The effect of torque on FA-I afferent responses. **A-C)** The proportion of FA-I afferents that showed responses scaled by torque based on (**A**) spike count, (**B**) peak discharge rate, and (**C**) first spike latency, with 2 N background normal force. **D)** The torque sensitivity (*S* ^all^) of each FA-I afferent in each torque phase and with three background normal forces shown in box and whisker plots. **E**) The torque sensitivity index comparing S ^low^ and S ^high^ for each afferent in each torque in box and whisker plots. See Figure 3 legend for detailed description.

During the unloading phase the pattern was similar, 59% (23/39) of FA-I afferents’ responses were scaled by torque (15 in clockwise, 23 in anticlockwise and 15 in both directions; **Figure 10A**). Eight FA-I afferents that had torque-scaled responses to one torque direction, had non-graded responses to the opposite torque direction, and two that weren’t scaled by torque at all had non-graded responses (*p* < 0.05, Mann-Whitney-Wilcoxon test).

##### Peak Discharge rate

The proportion of FA-I afferents that had peak discharge rates scaled in response to torque was lower compared to mean discharge rate scaling (**Figure 10A, B**). During the loading phase, 49% (19/39) of FA-I afferents had peak discharge rate scaling (14 in clockwise, 17 in anticlockwise and 12 in both directions). During the unloading phase, the peak discharge rate was scaled in 44% (17/39) of FA-I afferents (for 8 in clockwise, 15 in anticlockwise, and 6 in both directions).

##### First spike latency

First spike latencies were scaled by torques in 53% (21/39) of FA-I afferents (14 in clockwise, 15 in anticlockwise and 8 in both directions; **Figure 10C**). Thus, the number of afferents capable of encoding torque magnitude by means of spike timing is similar to that based on mean discharge rates. On average, FA-I afferent first spike latencies were 45.6% shorter in response to 7.5 mNm compared to 3.5 mNm torque.

#### FA-I torque sensitivity comparison

The analyses of torque sensitivity *S*_T_^all^ of FA1 afferents to both torque directions during loading and unloading phases, with 2 N normal force, revealed there was no difference in FA- I torque sensitivity between the loading and unloading phases (*p* = 0.17, *F*(1,194) = 1.9, two-way repeated measures ANOVA), and no difference between anticlockwise and clockwise torques (*p* < 0.11, *F*(1,194) = 2.5, two-way repeated measures ANOVA). The torque sensitivity increase index *I*_T_ values indicated that FA-I afferents were typically more sensitive to stimuli with higher torque magnitudes and concomitant faster torque increase rates (**Figure 10E**).

We found no difference in *I*_T_ between torque phases (W = 127, *p* = 0.80, Mann-Whitney-Wilcoxon test).

#### Effect of normal force on FA-I afferent torque responses

The background normal force appeared to influence FA-I afferent sensitivity to torque. **Figure 11A** shows the proportion of afferents that had mean discharge rates influenced by torque in at least one direction with different levels of background normal force. During the loading phase, the highest proportion of afferents with torque-scaled mean and peak discharge rates, and first spike latencies were found with 2 N background normal force, **Figure 11A-C**. Next, we investigated the relationship between background normal force and torque sensitivity for each FA-I afferent that was tested with all three normal forces (n = 13). In most FA-I afferents the background normal force scaled torque sensitivity 62% (8/13) (median correlation coefficient = -0.33, range = -0.99 – 0.25), and in all cases torque sensitivity decreased with increasing background normal force (Figure **11D**). Torque sensitivities and RFCs of FA-I afferents that were unaffected by changes in background normal force are shown in **Figure 11E**.

**Figure 11.**
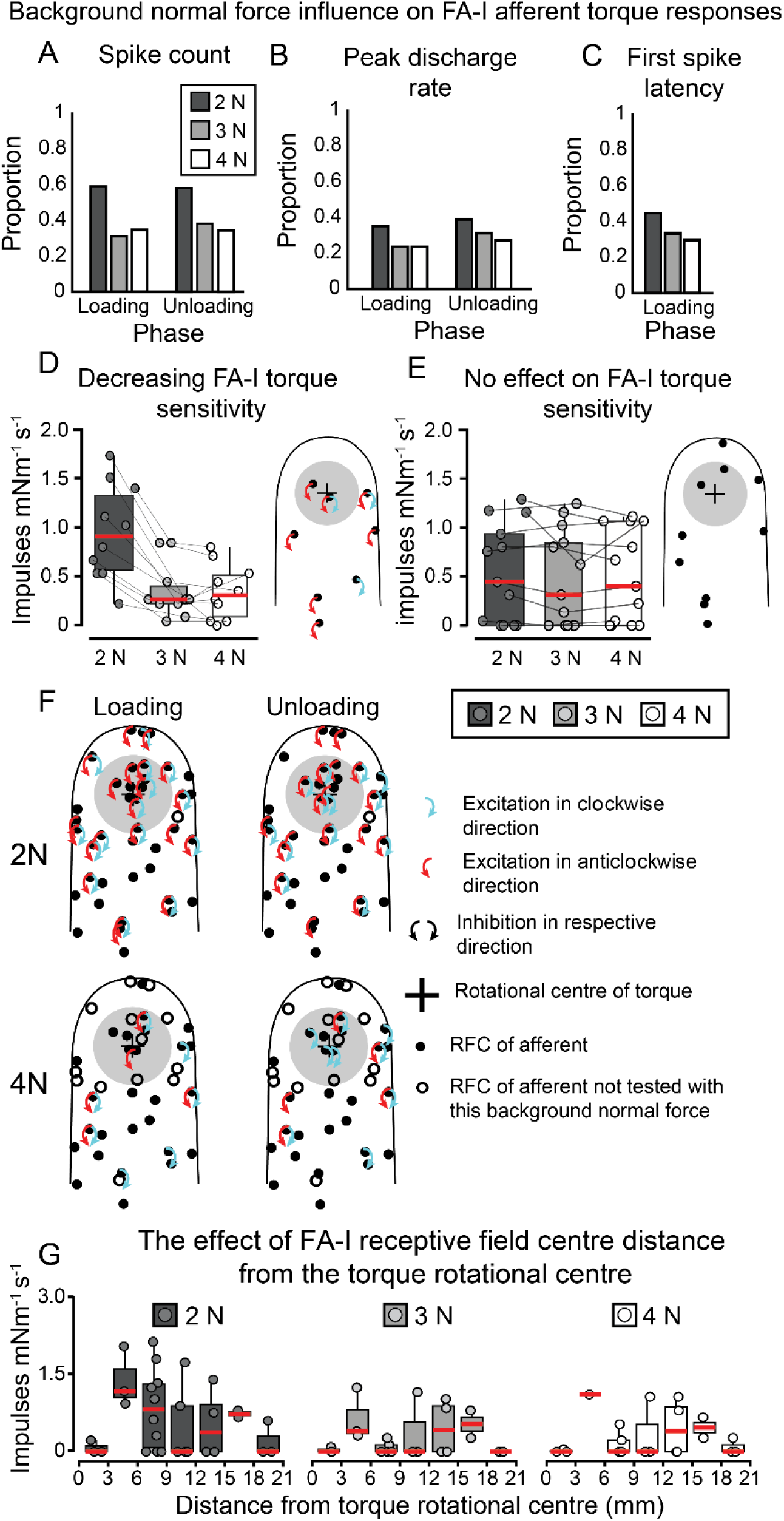
The effect of normal force and receptive field centre location on FA-I afferent’s torque responses. **A-C)** The proportion of afferents that were scaled by torque in at least one direction as measured by (**A**) spike count, (**B**) peak discharge rate, and (**C**) first spike latency for each normal force. All three metrics were calculated in the loading and unloading phases, as FA-I afferents were unresponsive to torques in the plateau phase. **D-E)** FA-I torque sensitivities with 2, 3, and 4 N background normal forces, separated by afferents that had significantly decreasing torque sensitivity with increasing background normal force **(D)**, or were unaffected by background normal force **(E). F)** The receptive field centre (RFC) location of each afferent and the direction of torque scaling are plotted for each torque phase across a standard fingertip. **G)** Sensitivity values from the torque loading phase are shown for FA-I afferents against their RFC distance from the stimulus rotational centre. FA-I afferents were mostly unresponsive to torque within 3 mm of the stimulus rotational centre. With 2 N background normal force, FA-I afferents showed the highest torque sensitivity when their RFCs were between 3-9 mm from the stimulus rotational centre. See Figure 5 legend for detailed description.

#### Relationship between receptive field location and torque sensitivity

The pool of FA-I afferents tested had RFCs distributed over most of the distal phalanx (**Figure 11F**). Like SA-I afferents, we expected to see torque sensitivity increase with FA-I RFC distances from the stimulus centre. **Figure 11G** shows the torque loading phase sensitivity (S_T_^all^) with three background normal forces. FA-I afferents were almost unresponsive to torque within 3 mm of the stimulus centre. Further from the stimulus centre, FA-I torque sensitivity increased between 3-6 mm, particularly with 2 N background normal force but their torque sensitivity varied more between 6-18 mm, before reducing to almost zero between 18-21 mm. Generally, the relationship between torque sensitivity and distance from the stimulus centre appeared to show an inverted U-shaped curve with minimal torque sensitivity at the stimulus centre and beyond 18 mm from the stimulus centre, which resembled that seen with SA-I afferents, but less consistent.

### Comparison of torque sensitivity and discriminability of human and monkey afferent types

It is well established that all tactile afferent types respond to most somatosensory stimuli of sufficient strength, but each afferent is well-adapted to encode some types of stimuli better than others. The stimuli used in the present study were too slow to excite FA-II afferents, but did reliably excite FA-I, SA-I and SA-II afferents. Previously it was demonstrated that FA-I and SA-I afferents in monkeys could encode torque magnitude, while SA-I afferents were better for encoding torque direction (Birznieks et al., 2010; Redmond et al., 2010a; Redmond et al., 2010b; Fu et al., 2012; Khamis et al., 2015). Here we sought to determine which human afferent types were better at discriminating torque magnitude and direction, and compare torque discriminability between human and monkey afferents, noting that SA-II afferents are absent from the fingers of monkeys.

#### Torque magnitude sensitivity

When comparing data obtained in the current study with Birznieks et al. (2010) it was generally observed that monkey SA-I afferents responded with greater sensitivity to torque magnitude changes than human SA-I afferents. Therefore, we sought to compare S_T_^all^ between humans and monkeys (human data previously shown in **Figure 7D**). We used a two-way repeated measures ANOVA to compare S ^all^ means and found a significant variation in the interaction between monkey and human S_T_^all^ in the torque loading and plateau phases (*p* = 0.067, *F*(1,198) = 14.3). Monkey SA-I mean S_T_^all^ in the torque loading phase was 1.20 ± 0.11 impulses mNm^-1^ s^-1^, which was nearly twice that of human afferents, 0.68 ± 0.08 impulses mNm^-1^ s^-1^ (*p* = 0.004, one-way post hoc test with Bonferroni adjustment) (**Figure 12A and B**). In the torque plateau phase monkey SA-I afferents had a mean S_T_^all^ of 2.00 ± 0.22 impulses mNm^-1^ s^-1^ which was more than five times greater than human afferent S_T_^all^ of 0.37 ± 0.05 impulses mNm^-1^ s^-1^ (*p* < 0.001 one-way post hoc test with Bonferroni adjustment). Similarly, investigation of FA-I afferent torque magnitude sensitivity found that monkey afferent S ^all^, 1.60 ± 0.18 impulses mNm^-1^ s^-1^, was significantly greater than human FA-I afferents, 0.77 ± 0.08 impulses mNm^-1^ s^-1^ in the torque loading phase (*p* < 0.001, Student’s t-test) (**Figure 12A and B**). Comparatively, human SA-II torque sensitivity averages in the loading (0.48 ± 0.07 impulses mNm^-1^ s^-1^) and plateau (0.46 ± 0.08 impulses mNm^-1^ s^-1^) phases were similar to human SA-I and FA-I afferents but were much lower than monkey SA-I and FA-I afferents.

**Figure 12.**
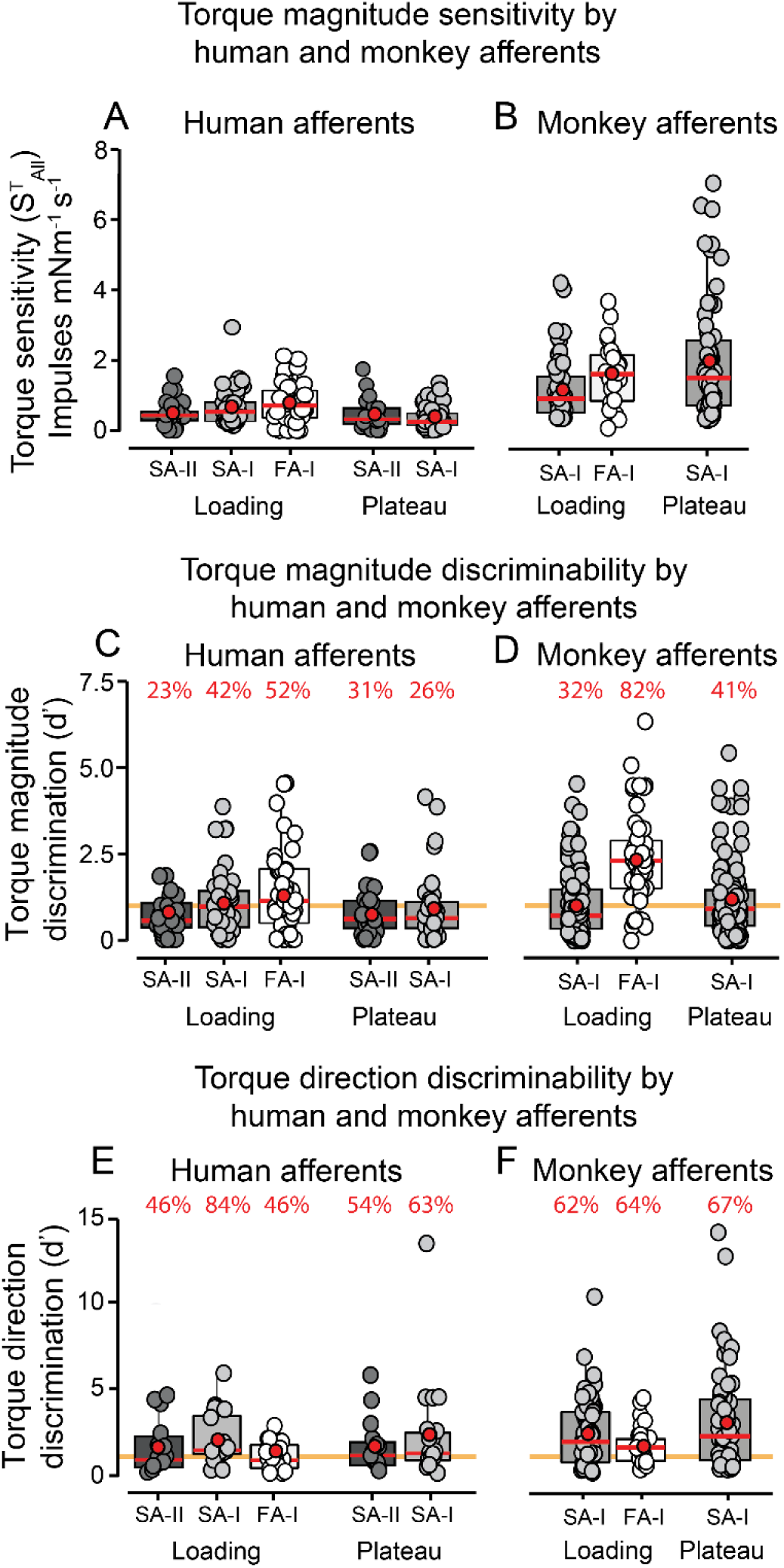
Comparison of human and monkey FA-I, SA-I, and SA-II torque magnitude and direction discriminability. **A-B)** Afferent torque sensitivity (*S*_T_^all^) of each afferent type in **(A)** humans and **(B)** monkeys. **C)** The torque magnitude discriminability of each human afferent type during the torque loading and plateau phases, with 2 N background normal force, in box and whisker plots. Each filled circle is the d’ value computed from the mean and standard deviation of the spike count in response to six torque trials with 3.5 mNm torque and 5.5 mNm torque, applied to a single afferent. Magnitudes were only compared between torques applied in the same direction, but the figure includes d’ values for both torque directions. **D)** Same as **C** but for monkey FA- I and SA-I afferent torque responses with 2.5 N background normal force. **E)** The torque loading and plateau phase direction discriminability of human SA-II, SA-I, and FA-I afferents with 2 N background normal force. d’ values were computed from the mean and standard deviation of spike counts in response to 7.5 mNm torque for clockwise and anticlockwise torques. **F)** Same as **E** but for monkey SA-I and FA-I afferents in response to 5.5 mNm torques, with 2.5 N background normal force. Red percentages above bars indicate the proportion of each afferent with d’ values greater than 1. Red horizontal lines and red circles indicate medians and means, respectively, and box boundaries indicate the upper and lower quartiles. Upper whisker shows the largest observation less than or equal to the upper box hinge + (1.5 × inter-quartile range), and the lower whisker shows the smallest observation greater than or equal to the lower box hinge – (1.5 × inter-quartile range). Gold coloured lines indicate the threshold d’ value of 1.

#### Torque magnitude discriminability

Human and monkey D-prime (d’) values were computed between the number of spikes evoked in response to 3.5 and 5.5 mNm torque during the torque loading and plateau phases for both torque directions, with 2.5 N background normal force (monkeys). Two d’ values were obtained for each afferent, one for each of the two torque directions. A d’ value of 1 was set as a threshold, such that if d’ was greater than 1 the two torque magnitudes were deemed discriminable from an afferent’s responses. About half of the human SA-I (42%) and FA-I (52%) afferent responses could discriminate torque magnitude in the loading phase (SA- I, 16/38 responses d’ > 1, median d’ = 0.9; FA-I, 25/48 responses d’ > 1, median d’ = 1.1) (**Figure 12C**), but the proportion was lower (23%) for SA-II afferents (6/26 responses d’ > 1; median d’ = 0.5). In the plateau phase SA-II afferent discriminability improved and 31% of afferents could discriminate torque (8/26 afferents with d’ > 1; median d’ = 0.53), while the proportion of SA-I afferents during plateau phase showed decreased discriminability of 26% (SA-I 10/38 responses d’ > 1, median d’ = 0.55) (**Figure 12C**). Compared to their human counterparts, monkey SA-I afferents were worse at discriminating torque magnitude in the loading phase, as a smaller 32% proportion (37/116 afferent responses with d’ > 1, median d’ = 0.67) of SA-I afferent responses was able to discriminate torque magnitude in the loading phase. The opposite was the case in the plateau phase, as 41% (47/116 afferent responses with d’ > 1, median d’ = 0.86), of monkey SA-I afferent responses could discriminate plateau torque magnitude, which was 15% more than the proportion of discriminable human SA-I afferent responses (**Figure 12C and D**). Monkey FA-I afferents appeared to be much better than their human counterparts in discriminating torque magnitude in the loading phase, with 82% of FA-I afferent responses able to discriminate torque magnitude (41/50 afferents with d’ > 1) and a median d’ of 2.3, which was more than double the human FA-I median d’ of 1.1 (**Figure 12C and D**).

#### Torque direction discriminability

Next, we compared torque direction discriminability among human and monkey afferent types by computing d’ from spike counts less background in response to both torque directions, of magnitude 5.5 mNm. In the loading phase 46% of both human SA-II and FA-I afferents (SA-II, 6/13 afferents with d’ > 1, median d’ = 0.91; FA-I, 11/24 afferents with d’ > 1; median = 0.88) could discriminate torque direction, but the proportion of discriminable SA-I afferents’ responses (84%) was nearly twice as large (16/19 afferents with d’ > 1; median d’ = 1.47) (**Figure 12E**). SA-II and SA-I discriminability was similar in the plateau phase, and there were no large differences between proportions of SA-I (63%, 12/19 afferents with d’ > 1; median = 1.3) and SA-II (54%, 7/13 afferents with d’ > 1; median = 1.2) afferents able to discriminate torque direction (**Figure 12E**).

Torque direction discriminability in monkey afferents was superior to their human counterparts in the torque loading phase for SA-I (62%, 36/58 afferents with d’ > 1; median 1.9) and FA-I (64%, 16/25 afferents with d’ > 1; median 1.5) afferents, and in the plateau phase for SA-I afferents (67%, 39/58 afferents with d’ > 1; median 2.2) (**Figure 12F**).

## Discussion

This study is the first to demonstrate how human SA-II, SA-I and FA-I afferents innervating the fingerpads respond to torque. Previous studies (Birznieks et al., 2010; Redmond et al., 2010a; Redmond et al., 2010b; Fu et al., 2012; Khamis et al., 2015) have used data obtained in monkeys, which despite possessing SA-II afferents in hairy skin, lack them in the glabrous skin of the hand (Johnson, 2001). We show that human SA-II afferents are highly sensitive to torque magnitude and direction and can readily encode these torque features. Moreover, SA- II, SA-I, and FA-I afferent responses could discriminate between different torque magnitudes and directions, but a population of the three afferent types are likely necessary to encode the full range of torque parameters. Monkey SA-I afferents were significantly more sensitive to torque magnitude than human afferents, whereas monkey FA-I afferents showed both greater torque magnitude sensitivity and superior discrimination of torque magnitude and direction.

### Single afferent responses to torque

Both the magnitude and direction of torque influenced FA-I, SA-I, and SA-II afferent activity in humans. During the loading phase, most (85%) SA-II afferents had mean discharge rates that were scaled by torque in at least one direction. Torques applied in either direction could enhance or supress SA-II afferents, which resulted in a diverse array of response behaviours to different torque magnitudes and directions that may, in part, derive from an SA-II afferent’s associated receptor (presumably the Ruffini ending) location relative to the rotational centre, and its anchoring within tissue. SA-II afferents are commonly found in or near the fingernail border (such as the afferent in **Figure 4A-C)** and might experience distinctive tangential force effects due to shear stress developing in the skin where the compliant tissue borders the stiff fingernail (Dandekar et al., 2003; Shimawaki and Sakai, 2007; Birznieks et al., 2009; Ho and Hirai, 2015; Wolterink et al., 2019).

### Human torque parameter sensitivity and discriminability

Encoding torque changes during the loading phase is critical to ensure grasp stability, but some dexterous tasks may also require maintenance of an object’s orientation and decreasing torque could indicate that an object is rotating and losing the desired orientation in the grip (Goodwin et al., 1998). SA-II afferents were more suitable to signal changes in torque magnitude during the plateau phase, corresponding to situations involving object orientation maintenance. By contrast, SA-I and FA-I afferents typically were more sensitive to dynamic changes in torque loading and unloading, as shown previously in macaques (Birznieks et al., 2010; Khamis et al., 2015).

All three afferent types could discriminate torque direction during torque loading. Eighty-three percent of torque-sensitive SA-II afferents appeared to have a torque directional preference, whereas only 54% of SA-I afferents and no FA-I afferents had obvious torque directional preferences, based on torque sensitivity data. However, d’ analyses showed that all three afferent types had discriminable responses between torque directions and there did not appear to be an afferent type that was best for encoding torque direction. This suggests that, the contribution of different individual afferents and afferent types signalling torque parameters changes depending on the task and stimulus parameters.

When fast signalling of torque magnitude is concerned, FA-I afferents potentially can deliver superior performance as the first spike latencies were scaled by torque in (58%) FA-I afferents, which was also the case in macaques (Birznieks et al., 2010). Therefore, FA-I afferents can rapidly signal torque changes precisely at onset, providing an advantage in controlling object orientation during object manipulation.

### Human and monkey afferent torque sensitivity and discriminability

We compared torque sensitivity and used d’ analysis to compare the discriminability of torque parameters by each afferent type in humans and monkeys. Remarkably, monkey afferents, particularly FA-Is, were significantly more sensitive to torque magnitude than human afferents. Human SA-Is showed decreased responsiveness and lower torque magnitude discrimination ability during the plateau phase in comparison to the loading phase, whereas monkey SA-Is performed better during the plateau phase. In both species, slowly adapting afferents were better suited to discriminate torque direction than FA-I afferents, and again monkey SA-I and FA-I afferents appeared to outperform their human counterparts. This demonstrates that input from all tested afferent types combined can provide the richest information about torque parameters.

An afferent information decoding model trained on macaque SA-I and FA-I afferent responses to normal force and torque was able to estimate instantaneous torque magnitude (Khamis et al., 2015). We found that torque magnitude discriminative capacity in humans was lower in FA-I afferents during the loading phase and in SA-Is during the plateau. SA-II afferents may, therefore, become a valuable addition to improve information content about torque. Sustained firing may be crucial to follow and make corrective motor responses when an object’s orientation should be maintained over longer periods without visual input (Johansson and Westling, 1987; Kinoshita et al., 1997). Due to similar sensitivity, one may wonder whether the nervous system would preferentially use one afferent type. We believe that in natural environments the challenge is to extract relevant information when multiple stimuli interact at the same time, which may be unlike the changes in constrained stimulus parameters in most studies. In such situations, combining inputs from a diverse afferent population might represent a computational advantage to disentangle the stimulus feature of interest (Khamis et al., 2015).

### Normal force effects and the spatial pattern of afferent torque responses

Macaque fingertips are approximately half the size of human fingertips, and typically have more bulged fingertip pulp tissues creating softer volar surfaces. Therefore, a larger proportion of the monkey fingertip is stimulated, despite using the same probe size in both studies, and they are more readily deformed by force and torque stimuli. Most human SA-I and FA-I afferents that were not influenced by torque had RFCs greater than 12 mm from the torque rotational centre, which suggests that torque stimulus effects did not reach their receptive fields.

During the loading phase, most afferents of each type had decreased torque sensitivity with larger background normal forces. Better response modulation to torque at lower normal forces might, in part, be caused by afferents responding to microslips arising close to the edge of the probe (Johansson and Westling, 1984; Westling and Johansson, 1987). In contrast, for other afferents—especially those with RFCs located outside the contact area—a larger normal force may cause a more profound fingertip deformation and thus a torque-generated skin deformation pattern could reach fingertip skin further away from the rotational centre. In the central ‘stuck’ zone, which doesn’t slip, FA-I afferents did not reliably respond to torque (André et al., 2011; Adams et al., 2013; Delhaye et al., 2014; Delhaye et al., 2021; Schiltz et al., 2022). Nearer the edge of the contact area, and just outside it, both FA-I and SA-I afferents were typically more sensitive to torque and most likely responded to changes in tangential or shear stress in the skin as well slip events, which may also be reflected in the instantaneous discharge rate profile irregularities of the of SA-II afferents observed in our study.

Although we did not systematically change the stimulus friction, it is expected that the contact area size would depend on frictional conditions and thus would influence the way afferents respond to torque. This is supported by previous studies suggesting that torque information is encoded while superimposed on a range of normal forces (Birznieks et al., 2010; Khamis et al., 2015), in which SA-I and FA-I afferents are generally more sensitive to torque with lower normal force when peripheral slips are more likely to occur. We used a high friction surface which, in combination with higher background normal force, would limit partial slip probability under the stimulation probe. With more slippery surfaces, the slipping area with the same contact force will be larger, but tensile strain caused by rotational movement will be smaller. Thus, it is expected that the input from SA-II afferents would decrease while it would become more prominent from FA-I and SA-I afferents within the slipping zone under the probe. We hypothesise that during natural manipulative tasks small rotational slips are a significant sensory source of frictional information, as torques are almost always present during gripped object handling and are permitted to occur without endangering grip safety. Therefore, future studies should investigate frictional effects and torque effects in combination with linear tangential force mimicking object manipulation.

## Conclusions

For the first time, we have demonstrated that human SA-II afferents encode torque magnitude and direction with high sensitivity. Interestingly monkey afferents were generally superior in their sensitivity and discrimination of torque parameters. SA-II afferents may increase the richness of information about torque stimuli that populations of human tactile afferents provide to the sensorimotor system during natural manipulations. This is particularly important as human SA-I afferents’ ability to differentiate torque parameters appear to be inferior to monkeys, especially during the plateau phase. Finally, SA-II afferents could contribute unique inputs to the nervous system, providing computational advantages to disentangle and extract information about multiple concurrently changing tactile stimuli during natural object manipulation.

## Acknowledgements

We thank Prof Antony W Goodwin for contribution to the inception of this study as well designing and providing stimulators from his laboratory. We thank Dr Rachael Brown for help with spike sorting procedures. This project was funded by the Australian Research Council (ARC).

